# Cryo-EM structure of a methanogen nitrogenase-PII protein supercomplex

**DOI:** 10.1101/2025.09.09.675011

**Authors:** Rajnandani Kashyap, Thomas M. Deere, Ahmed Dhamad, Melissa Chanderban, Monika Tokmina-Lukaszewska, Brian Bothner, Daniel J. Lessner, Edwin Antony

**Affiliations:** Department of Biochemistry and Molecular Biology, St. Louis University School of Medicine, St. Louis, MO, USA; Department of Biological Sciences, University of Arkansas, Fayetteville, AR, USA; Department of Chemistry and Biochemistry, Montana State University, Bozeman, MT, USA

**Author notes:** Department of Biological Sciences, Wasit University, Wasit, Iraq (Ahmed Dhamad).

## Abstract

Nitrogenases are metalloenzymes that catalyze the reduction of atmospheric dinitrogen to ammonia, sustaining the global nitrogen cycle. While bacterial nitrogenase has been extensively characterized, the architecture and regulation of archaeal nitrogenases remain unknown despite longstanding evidence of nitrogen fixation in methanogens. Here we report a 3.2 Å cryo-electron microscopy structure of a native nitrogenase–PII protein supercomplex from *Methanosarcina acetivorans*. The structure reveals an unprecedented assembly of three NifDK heterotetramers bridged by six NifI_1,2_ heterotrimeric PII complexes, which sterically block NifH association and lock the enzyme in an inactive state. The PII complexes display asymmetric binding of ADP and 2-oxoglutarate, coupling nitrogenase inhibition directly to cellular energy and nitrogen status. Addition of 2-oxoglutarate and ATP releases the NifI complexes, stimulating a threefold increase in NifDK activity *in vitro*. This higher-order architecture uncovers a previously unrecognized regulatory strategy in methanogens, in which PII proteins drive nitrogenase oligomerization to control activity. The discovery that nitrogenase activity may be modulated through direct assembly into higher-order structures opens new avenues for exploring nitrogenase evolution, regulation, and biotechnological applications.

**One sentence summary:** Discovery of a nitrogenase-PII protein supercomplex in methanogens, uncovering a metabolite-gated assembly mechanism for nitrogenase inhibition.

## INTRODUCTION

Biological nitrogen fixation (diazotrophy) is a critical process in the global nitrogen cycle, yet it is confined to a limited group of bacteria and archaea. All known and predicted diazotrophs rely on molybdenum (Mo)-dependent nitrogenase to catalyze the energetically demanding reduction of atmospheric dinitrogen (N_2_) to ammonia (NH_3_), following the stoichiometry^1,2^:

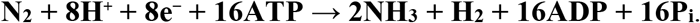

Mo-nitrogenase is a highly oxygen-sensitive metalloenzyme composed of two components: dinitrogenase (a NifD-NifK heterotetramer, also known as the MoFe protein) and dinitrogenase reductase (a NifH homodimer, or Fe protein). NifH contains a [4Fe-4S] cluster that transfers electrons to the P-cluster (8Fe-7S) of NifDK, which in turn delivers electrons to the FeMo-cofactor (FeMo-co or M-cluster), the active site for N_2_ reduction^3-5^. Transfer of one electron by NifH requires the hydrolysis of two ATP molecules^2,6,7^. Given the high energy cost, nitrogenase expression and activity are tightly regulated at multiple levels - transcriptional, translational, and post-translational - and are closely integrated with cellular carbon metabolism and energy status^8-10^.

PII signal transduction proteins are central regulators of nitrogen fixation and assimilation in diazotrophs^11^. These ancient and highly conserved proteins are found across all domains of life^12^. Canonical PII proteins (e.g., GlnK, GlnB, GlnZ) form homotrimers, each with three ligand-binding sites located at the interface between subunits. Each monomer adopts a ferredoxin-like α/β fold with three characteristic loops (T, B, and C). Ligand-induced conformational changes in the T-loop are key to modulating interactions with target proteins. PII proteins sense the cellular carbon and nitrogen status by binding 2-oxoglutarate (2OG), a TCA cycle intermediate and universal indicator of nitrogen limitation^13^. They also bind ADP and ATP to monitor the cell’s energy state. These ligand interactions influence PII binding to various targets, including enzymes, transporters, and regulatory proteins^14^.

PII proteins are known to indirectly control the activity of nitrogenase in diverse bacteria, including anaerobes, aerobes, and oxygenic phototrophs. The best example is the DraT-DraG system, where under nitrogen-replete conditions the PII protein GlnB activates DraT, which reversibly inactivates NifH via ADP-ribosylation, thereby preventing association with NifDK. This modification is triggered by low intracellular 2OG, signaling nitrogen sufficiency^11^.

In contrast, nitrogenase in archaea is restricted to anaerobic methanogens and methanotrophs. Methanogens, among the most ancient lineages of life, are thought to be the progenitors of nitrogenase^15,16^. They typically possess a minimal *nif* gene cluster (*nifHI*_*1*_*I*_*2*_*DKEN*), encoding the catalytic components (NifHDK), the FeMo-co assembly scaffold (NifEN), and two PII proteins (NifI_1_ and NifI_2_)^1,16^. Methanogens operate near the thermodynamic limits of life, conserving significantly less energy during methane production than aerobic diazotrophs like *Azotobacter vinelandii* or cyanobacteria<u>^17^</u>. Thus, efficient regulation of nitrogenase is vital for survival under fluctuating nitrogen conditions.

In studied methanogens - including *Methanococcus maripaludis, Methanosarcina barkeri, Methanosarcina mazei*, and more recently *Methanosarcina acetivorans* - nitrogenase production is regulated by fixed nitrogen (e.g., ammonia) availability^18-24^. Unlike proteobacteria, methanogens control nitrogenase activity through direct interaction of the NifI_1,2_ complex with NifDK. Studies with *M. maripaludis* have shown that under nitrogen-replete conditions, the heteromeric NifI_1,2_ complex binds NifDK, likely blocking electron transfer from NifH and thereby inactivating nitrogenase. The presence of 2OG and ATP promotes dissociation of this complex, restoring activity^22,25,26^. Notably, NifI_1,2_ is the only known heteromeric PII complex and the only PII variant that directly interacts with nitrogenase.

Here, we demonstrate that in *M. acetivorans*, the NifI_1,2_ complex forms a heterotrimer composed of two NifI_1_ subunits and one NifI_2_ subunit. Six NifI_1,2_ complexes assemble into a novel 30 subunit ∼898 kDa supercomplex by binding six NifDK heterodimers. Importantly, this interaction not only prevents NifH from accessing NifDK but also likely induces oligomerization of NifDK itself, forming a unique inhibitory structure. These findings reveal a previously unrecognized mechanism of nitrogenase regulation in methanogens, mediated by PII protein-dependent oligomerization.

## RESULTS

### *M. acetivorans* strain DJL70 expresses functional Strep-NifD that co-purifies with NifK, NifI_1_ and NifI_2_

To facilitate the purification of NifDK (MoFe protein) from *M. acetivorans*, strain DJL70 was generated using the CRISPR-Cas9 system to replace native *nifD* in parent strain WWM73 with *nifD* encoding a tandem Strep tag (*strep-nifD*) (**Fig. 1A**). Importantly, strain DJL70 grows similar to strain WWM73 when using N_2_ as the sole N source, indicating that Strep-NifD does not alter Mo-nitrogenase expression and activity (**Fig. 1B**). Also, like strain WWM73, and consistent with previous results^18,19^, Strep-NifD is only detected by Western blot in DJL70 cells grown without NH_4_Cl (**Fig. 1A**). Affinity purification of Strep-NifD results in the co-purification of NifK and NifI_1_ and NifI_2_, as seen by SDS-PAGE (**Fig. 1C**) and confirmed by mass spectrometry (Source Data File). In *M. maripaludis*, NifI_1_ and NifI_2_ form an inhibitory complex that binds NifDK to block N_2_ reduction in response to depletion of 2OG, an indicator of fixed N sufficiency^26^. Despite strain DJL70 grown only with N_2_ as the sole N source, a significant portion of NifDK appears in the NifI_1_-NifI_2_ (NifI_1_I_2_) bound inhibited state, similar to results obtained with *M. maripaludis*^25^. Importantly, the addition of 2OG was shown to the release of NifI_1_I_2_ from NifDK, allowing electron transfer from NifH thereby facilitating the reduction of N_2_ by NifDK^25,26^. To ascertain if co-purified NifI_1_I_2_ performs the same function in *M. acetivorans*, the acetylene reduction activity of purified NifDK was assayed with purified recombinant NifH (**Supplemental Fig. 1**) in reaction mixtures with or without 2OG (**Fig. 1D**). An approximate 2-fold increase in the specific activity (nmol ethylene reduced min^-1^ mg^-1^) of NifDK with 2OG was observed [537 ± 97 (-2OG) vs 1239 ± 67 (+2OG)], indicating that a significant portion of *in vivo* NifDK is in the inactive NifI_1_I_2_ bound inhibited state. To understand how NifI_1_I_2_ binds to NifDK, the affinity-purified complex was directly analyzed by single-particle cryo-EM without the addition of 2OG.

**Figure 1.**
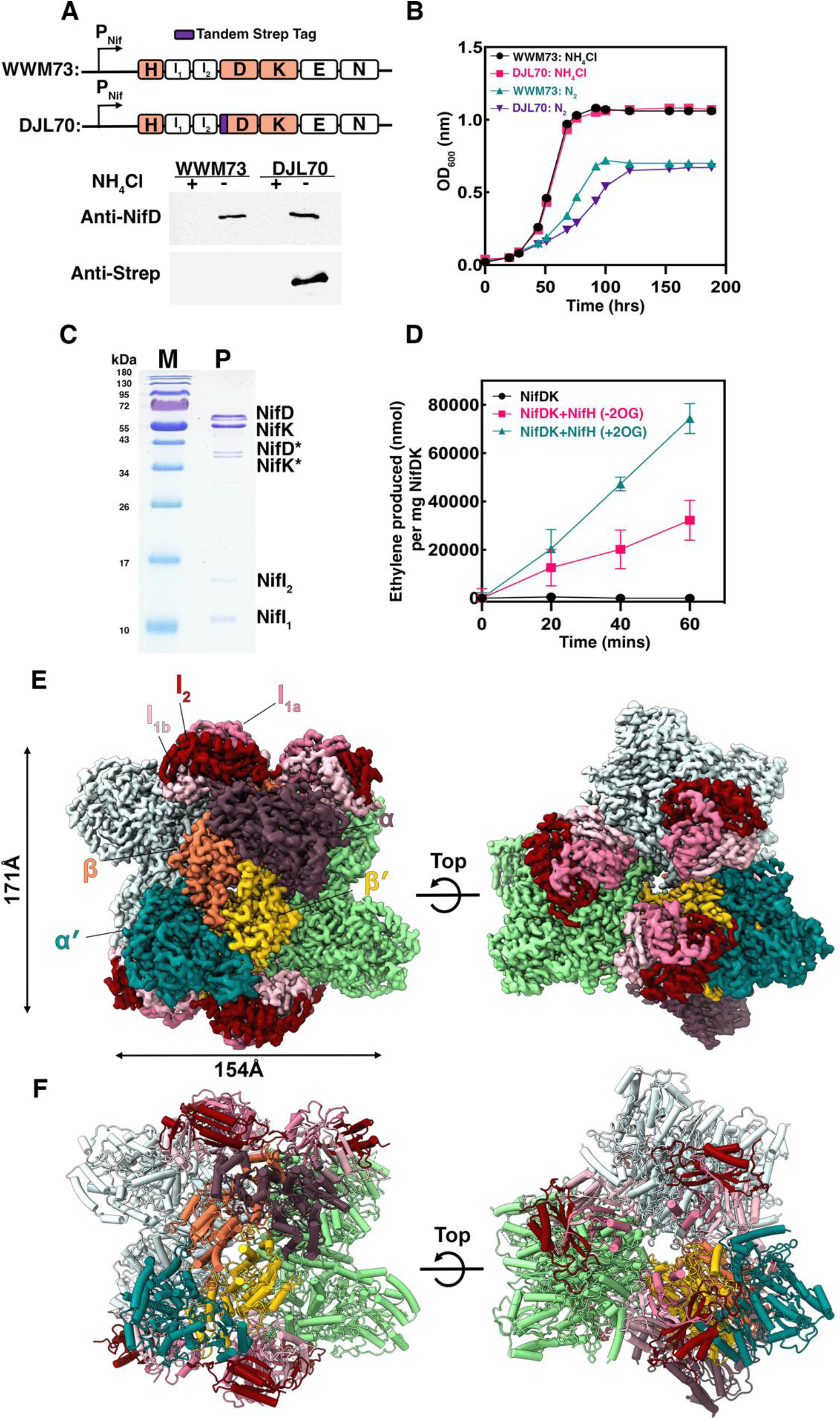
Genetic, biochemical, and structural analysis of nitrogenase function in *M. acetivorans*. **A**. Schematic comparison of the nifHI_1_I_2_DKEN operon in strains WWM73 and DJL70. A tandem N-terminal Strep-tag was introduced at the *nifD* locus in WWM73 using CRISPR-Cas9 to generate strain DJL70. The sub-panel below shows Western blot analysis of lysates from cells grown in **(B)**, probed with antibodies against NifD and the Strep tag. **B**. Growth of each strain in HS medium with methanol (125 mM) with either NH_4_Cl or N_2_ as the nitrogen source. **C**. SDS-PAGE analysis of a representative purification of Strep-tagged NifD, showing co-elution of NifK, NifI_1_, and NifI_2_ proteins, from strain DJL70, as described in the Materials and Methods. Two additional bands below the full-length NifD and NifK, marked with asterisks (*), were later identified as truncated forms of NifD and NifK. **D**. Effect of 2-oxoglutarate (2OG) on acetylene reduction activity of purified NifDK. Ethylene production was measured per mg of NifDK (75 µg) in the presence of recombinant NifH (150 µg), with or without 10 mM 2OG. Reactions without added NifH were included as a negative control. Data represent the mean±SD from three biological replicates. **E**. Representative Cryo-EM map of the nitrogenase super-complex resolved at 3.16 Å, revealing the overall architecture and spatial organization of its core components. The supercomplex consists of three NifDK units, each adopting an α_2_β_2_ architecture and coordinated by a NifI heterotrimer positioned on either side of the tetrameric assembly. NifD (α) and NifD′ (α′) are shown in eggplant and teal, while NifK (β) and NifK′ (β′) are colored coral and gold, respectively in one NifDK unit. The other two units are colored green and grey, respectively. The associated inhibitor proteins-NifI_1a_, NifI_1b_, and NifI_2_ are depicted in pale violet-red, pink, and maroon, respectively. **F**. Cryo-EM structure of the supercomplex highlighting a single NifDK tetrameric unit bound by two heterotrimeric NifI complexes on either end (colors are the same as in panel E).

### *M. acetivorans* NifDK forms a supercomplex with NifI_1_-NifI_2_

2D-classification of the cryo-EM images revealed the presence of four distinct particle populations in the dataset (**Supplemental Figs. 2, 3, & 4**). The largest volume showcased a structurally homogeneous complex which resulted in a high-resolution structure of the *M. acetivorans* NifDK-NifI_1_-NifI_2_ supercomplex at 3.16 Å (**Fig. 1E, F; Supplemental Table 1**). The supercomplex reveals a non-canonical assembly of nitrogenase comprising three NifDK heterotetramers, each associated with two heterotrimeric PII-inhibitor complexes consisting of two NifI_1_ (NifI_1a_ and NifI_1b_) subunits and one NifI_2_ subunit. For clarity in presentation, we here define each NifDK tetramer along with the associated NifI_1a,b_-NifI_2_ complexes bound on either half as one unit (the two NifI1 subunits are labeled a & b, respectively). Three such units form the supercomplex. In addition, we also define the two halves within each unit: For example, NifD (α) and NifK (β) subunits in unit 1 are labeled as α1, β1 and α1′, β1′. The NifI proteins bound in unit 1 are correspondingly labeled as NifI_1a_, NifI_1b_, NifI_2_, and NifI_1a′_, NifI_1b′_, NifI_2′_, respectively (**Fig. 2A**).

**Figure 2.**
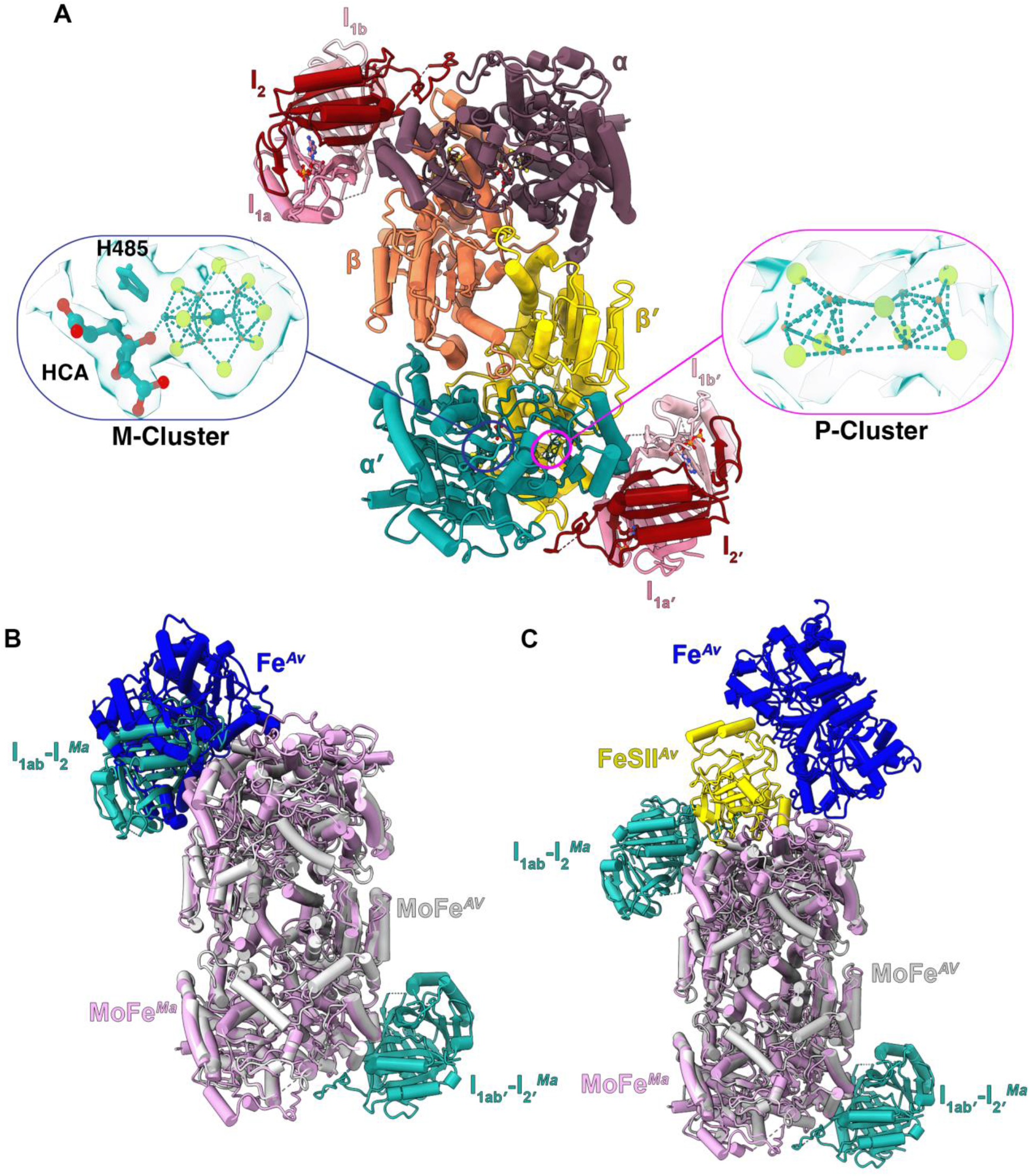
Structural features of individual supercomplex units and overlay with *A. vinelandii* NifDK bound to NifH and FeSII. **A**. Cryo-EM structure of an individual *M. acetivorans* (Ma) NifDK tetramer highlighting the core components of a single unit. Each unit adopts an α_2_β_2_ architecture to form the core NifDK complex and is bound by two NifI heterotrimers positioned on either side of the tetrameric assembly. In the NifDK unit, NifD (α) and NifD′ (α′) are shown in eggplant and teal, while NifK (β) and NifK′ (β′) are shown in coral and gold, respectively. The associated inhibitor proteins NifI_1a_, NifI_1b_, and NifI_2_ are depicted in pale violet-red, pink, and maroon, respectively. The callouts highlight the P-cluster, the M-cluster containing homocitrate (HCA), and the His-485 ligand. Each cluster is displayed with its corresponding carved density map. **B**. Overlay of a single Ma NifI (teal)–NifDK (pink) complex (Unit 1) from our cryo-EM structure with the asymmetric *A. vinelandii* nitrogenase NifDK (MoFe protein; grey; PDB: 7UT8) bound to NifH (Fe protein; blue). **C**. Overlay of a single Ma NifI (teal)–NifDK (pink) complex (Unit 1) from the supercomplex onto the *A. vinelandii* NifDK (grey) tetramer (MoFe protein; PDB: 8RHP) bound to FeSII (yellow) and NifH (Fe protein; blue).

Despite the unusual higher-order assembly, the core architecture of each NifDK unit remains conserved and resembles that of the well-characterized *Azotobacter vinelandii* (Av) nitrogenase (**Fig. 2A & B**). The NifD and NifK subunits form an α_2_β_2_ heterotetramer (MoFe-protein) and each half houses a M-cluster ([Mo-Fe_7_S_9_C•homocitrate]) and P-cluster ([Fe_8_S_7_]) (**Fig. 2A and Supplemental Figs. 5 & 6**). The M-cluster in *M. acetivorans* NifDK is coordinated by His-485, and Cys-267 from NifD while Lys-469 interacts with homocitrate (HCA) (**Supplemental Fig. 5A**). The P-cluster is coordinated by Cys-48, Cys-106, Cys-23, and Ser-141 from NifK along with Cys-58, Cys-84, and Cys-149 from NifD (**Supplemental Fig. 6B**). These contacts are identical in each NifDK half in all three units observed in the supercomplex. While the overall structure of NifDK in all three units are similar, minor structural changes are observed, especially around the M-cluster. His-485, positioned near the M-cluster, adopts different conformations in the NifDK halves in units 1 & 2 of the supercomplex (**Supplemental Fig. 6**).

### The PII heterotrimeric inhibitor complex coordinates assembly of multiple MoFe-proteins in the supercomplex

PII proteins are a conserved group of nitrogen-regulatory proteins that form homotrimeric complexes in structural studies^12,27^. In contrast, in the supercomplex, the NifI proteins form a heterotrimer (PII complex) with one subunit of NifI_2_ wedged between two copies of NifI_1_ (NifI_1a &_ NifI_1b_; **Fig. 2A**). Each NifDK half in the tetramer is bound to a PII complex. Comparison of the *A. vinelandii* (*Av*) nitrogenase and the PII-bound NifDK complex from *M. acetivorans* shows that the NifH and PII complex binding interfaces on NifDK overlap (**Fig. 2B**) and reveal how nitrogenase activity is switched-off by PII complex binding in methanogens. Recently, *Av* nitrogenase was shown to form inactive and oxygen-resistant assemblies promoted by Shethna protein II (FeSII)^28^. Like the PII complex in methanogens, FeSII is an inhibitor of nitrogenase in *Av* and drives the assembly of NifDK, along with NifH, into higher order complexes ^28^. Consistent with different functions, the binding of FeSII and the PII complex to the MoFe protein are distinct (**Fig. 2C**). The assembly of the supercomplex in *M. acetivorans* involves direct MoFe protein interaction, whereas Av MoFe proteins do not interact in higher order structures. These differences highlight the evolutionary divergence in regulatory strategies that involve oligomerization to inactivate nitrogenase in response to nitrogen sufficiency by anaerobic methanogens and to protect nitrogenase from oxygen in aerobic bacteria.

The nitrogenase supercomplex from *M. acetivorans* reveals extensive inter-subunit interactions among neighboring NifDK heterotetramers that contribute to the stability of the higher-order assembly (**Fig. 3**). The three NifDK complexes are arranged such that each tetramer is tilted ∼48º with respect to each other. The buried surface area is ∼1000 Å^2^ between adjacent NifDK tetramers in the supercomplex. An array of charged and polar residues forms a dense network of acid-base salt bridges and hydrogen bonds between the NifD and NifK subunits from adjacent units, with majority of the interactions mediated by NifK residues (**Fig. 3**). The spatial distribution of these interactions is symmetrical and therefore forms a cooperative electrostatic interface that stabilizes the trimeric arrangement. This supercomplex of *Ma* nitrogenase is distinct from other structures of bacterial nitrogenases that are typically observed as single NifDK heterotetramers^29^. There is no other reported evidence for such direct interactions between NifDK tetramers either under *in vivo* or *in vitro* conditions. Nitrogenase from aerobic bacteria have recently been shown to form a filamentous higher order structure, but this complex is formed through protein-protein interactions between nitrogenase and regulatory factors (e.g., FeSII)^28^. The presence of extensive electrostatic interfaces in *Ma* NifDK suggests a distinct regulatory mechanism unique to NifI-containing anaerobes. The stabilization of the trimeric assembly through inter-subunit salt bridges could lock the enzyme in a catalytically inactive state, representing a previously unobserved strategy for nitrogenase inhibition and assembly regulation. Interestingly, a NifH homodimer can be manually docked on NifDK within the supercomplex architecture with no steric hinderance if the NifI complex is removed (**Supplemental Fig. 7**).

**Figure 3.**
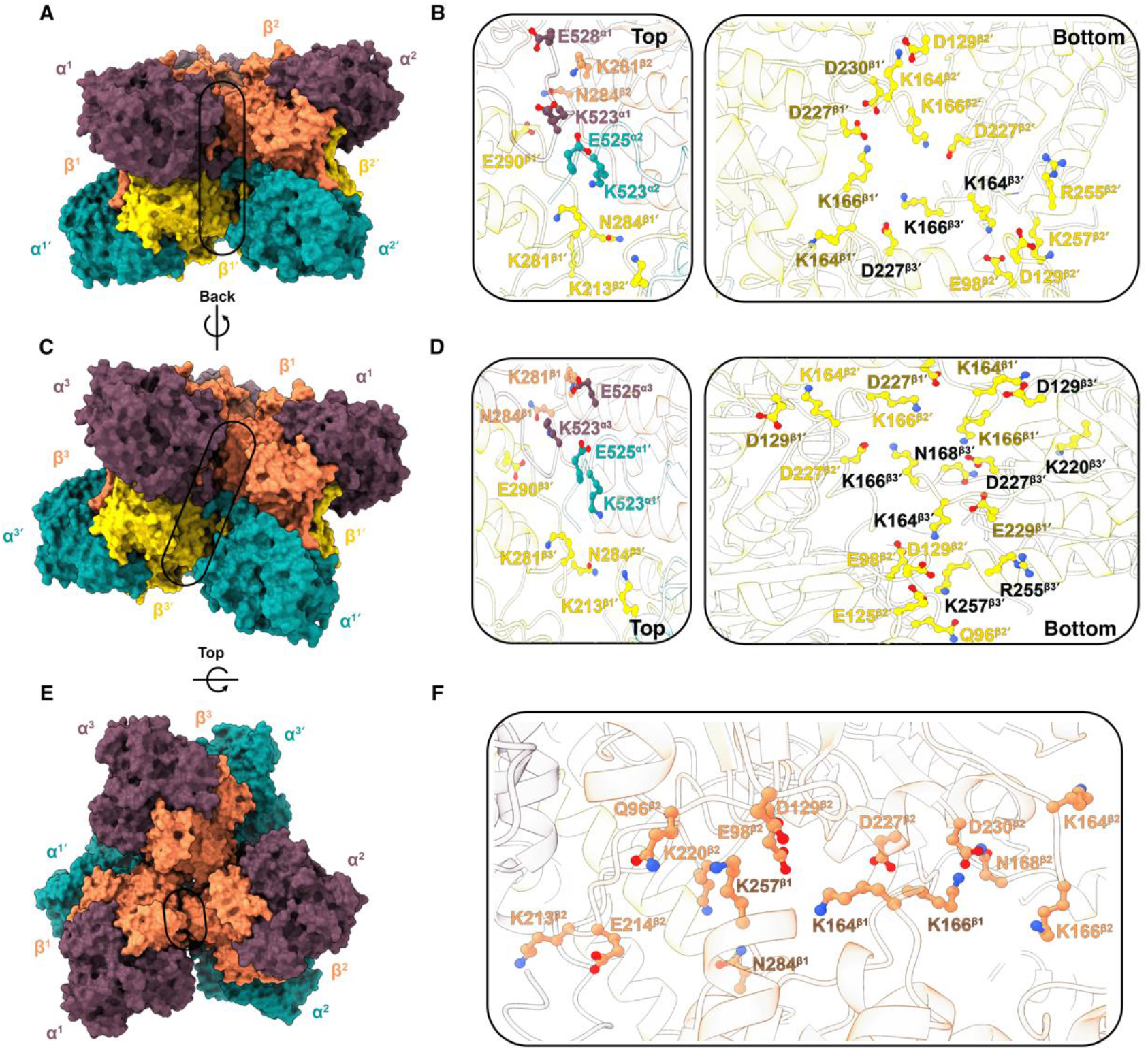
Interactions between NifDK tetrameric units within the supercomplex. **A**. Surface representation of NifDK subunits in the supercomplex. The interaction interface between Unit 1 and Unit 2 is highlighted. **B**. Close-up view of residues mediating this interaction. **C**. The interface between the NifDK subunits in Unit 1 and Unit 3 is indicated. **D**. Detailed view of residues involved in this interaction. **E**. Top view of the NifDK tetrameric units within the supercomplex, showing extensive interactions between the NifK (β) subunits. **F**. Enlarged view highlighting the individual residues involved in these interactions. In this figure, NifD (α), NifD′ (α′), NifK (β), and NifK′ (β′) are colored eggplant, teal, coral, and gold, respectively.

Each PII complex inhibitor in our structure is arranged as a NifI_1a_-NifI_2_-NifI_1b_ complex and is positioned to mediate interactions with two adjacent NifDK units in the supercomplex. An example of the NifI complex connecting two distinct NifDK units is shown in **Fig. 4**. This spatial configuration facilitates extensive inter-subunit contacts within the PII complex, predominantly mediated by salt bridges, with additional contributions from hydrogen bonding and hydrophobic interactions (**Fig. 4C-E)**. NifI_2_ interacts in *cis* with NifDK in the primary unit (**Fig. 4D**). In contrast, NifI_1a_-NifI_1b_ make extensive contacts in *trans* with NifDK of the adjacent unit (**Fig. 4C**). The PII complex and NifDK interactions are largely conserved across all three units, albeit with some differences in the total buried surface area within the interaction interface (**Supplemental Fig. 8**). Thus, in addition to extensive interactions between NifDK tetramers within the three units, each NifI complex also serves to connect and stabilize the three units within supercomplex.

**Figure 4.**
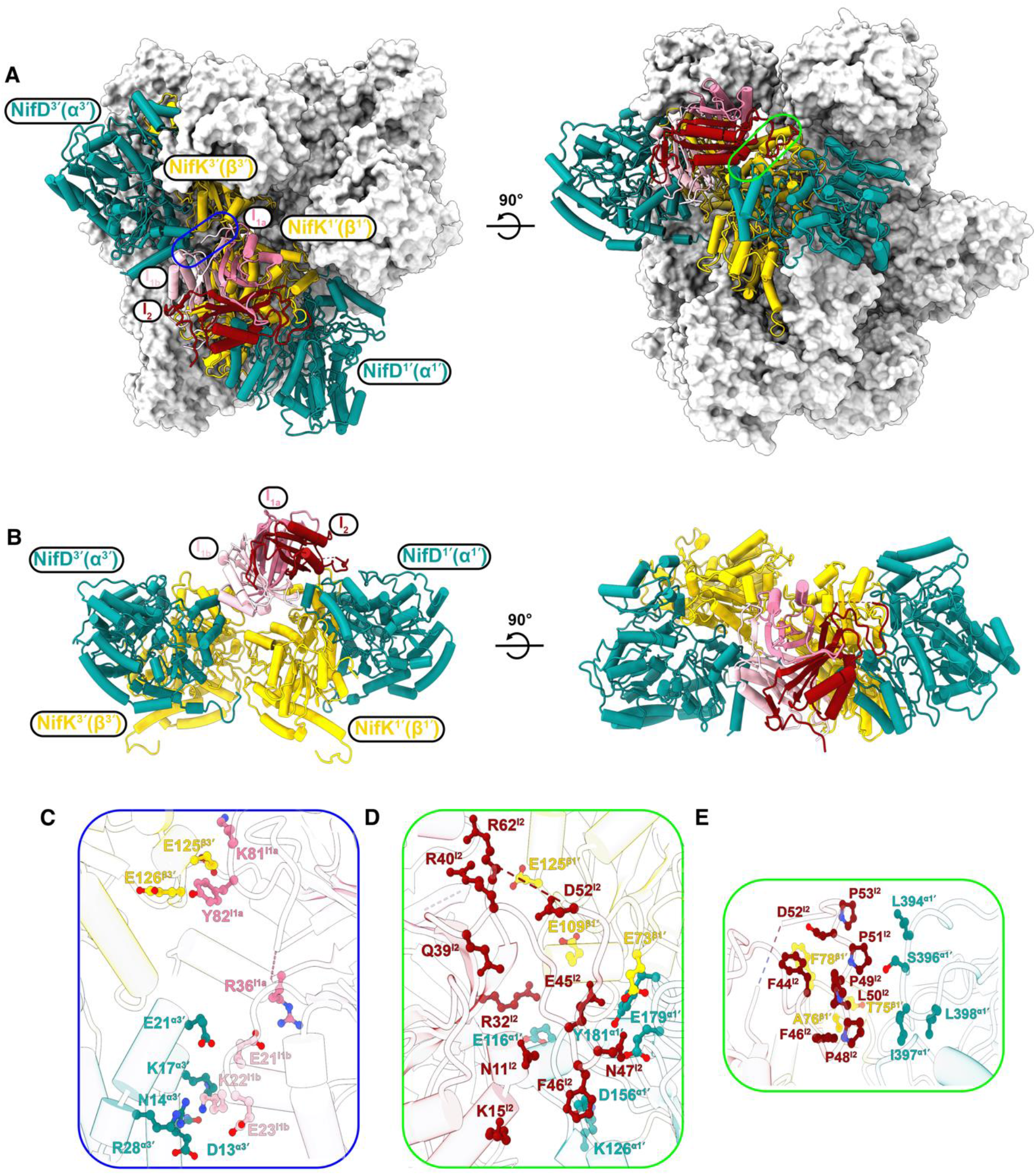
Interactions between the NifDK tetrameric unit and the NifI complex within the nitrogenase supercomplex. **A**. Cartoon representation of a single NifI complex interacting with NifDK tetramers of Unit 1 and Unit 3 and overlaid on the surface-rendered supercomplex. Interaction interfaces with Unit 1 and Unit 3 are highlighted by the blue and green demarcations, respectively. **B**. Close-up view of the NifDK units and associated NifI complex, illustrating their spatial arrangement and interface architecture. **C**. Detailed view of residues mediating the interaction between NifI_1a_ and NifI_1b_ of the NifI complex with NifDK of Unit 3. **D. & E**. Detailed views of residues involved in the interaction between NifI_2_ and NifDK. NifD (α), NifD′ (α′), NifK (β), and NifK′ (β′) are colored eggplant, teal, coral, and gold, respectively. The NifI subunits NifI_1a_, NifI_1b_, and NifI_2_ are colored pale violet-red, pink, and maroon, respectively.

### The NifI heterotrimeric complex is structurally similar to canonical homotrimeric PII proteins

Canonical PII proteins are homotrimers, consisting of monomeric units each typically 11-18 kDa. Each monomer exhibits a ferredoxin-like fold with three protruding flexible loops (T, B, and C; **Supplemental Fig. 9**). In the trimeric state, ligand binding sites are formed at the dimeric interface of two subunits, whereby the B-loop and the base of the T-loop of one subunit and the C-loop of the adjacent subunit form a pocket capable of binding ADP, ATP, and/or 2OG^12,14^ (**Supplemental Fig. 10**). As such, the trimer has three intercommunicating ligand binding sites. For example, GlnK in bacteria and archaea controls ammonium transport by the Amt transporter. At low intracellular levels of 2OG, which indicates ammonium sufficiency, GlnK binds ADP which results in the T-loop having a rigid extended confirmation that specifically binds within the channel of Amt effectively blocking ammonium transport. When ammonium is depleted, the intracellular levels of 2OG rise and GlnK subsequently binds 2OG along with ATP, resulting in the T-loop changing conformation and disassociation of GlnK from Amt^30,31^. Thus, the ADP-bound form of PII proteins typically binds target proteins and the ATP/2OG bound form typically does not.

NifI_1_ (11.8 kDa) and NifI_2_ (13.9 kDa) are similar in size to canonical PII proteins and contain the conserved residues involved in binding ADP/ATP and 2OG (**Supplemental Fig. 11**). A distinguishing difference is that NifI_2_ has a longer T-loop and an extended C-loop compared to NifI_1_. Importantly, both the T-loops of NifI_1_ and NifI_2_ were shown to be required for NifI_1_-I_2_ association with NifDK in *M. maripaludis*, consistent with the T-loops modulating target protein interaction^25^. Interestingly, the T-loops of NifI_1_ from both methanogens are quite similar in sequence, whereas the longer NifI_2_ T-loop sequences a markedly different, indicating there are likely differences in target protein specificity (**Supplemental Fig. 11**). Consistent with other canonical homotrimeric PII proteins, the NifI heterotrimer complex exhibits a ferredoxin-like fold, with the trimerization interface stabilized by a central β-sheet core as shown for the NifI complex bound to unit 1 in the supercomplex (**Fig. 5**). The T-loops for both NifI_1a_ and NifI_2_ are structured and extended, whereas the T-loop for NifI_1b_ is unresolved/flexible (**Fig. 5B**). Several residue-specific interactions reinforce the stability of the complex, including salt bridges and hydrophobic interactions (**Fig. 5C**). Collectively, these interactions promote formation of a stable heterotrimeric NifI complex. The NifI_2_ monomeric unit is identical in structure to that of GlnK, (**Supplemental Fig. 10A**), except for the T-loop and extended C-loop. All six NifI complexes are largely identical when superimposed (**Supplemental Fig. 12)**, with only differences in the T-loop conformation observed. Thus, although the NifI complex is composed of two separate proteins, the assembled heterotrimeric complex is indistinguishable to canonical PII homotrimers, indicating the NifI T-loop confirmation is also responsive to 2OG and ATP/ADP binding.

**Figure 5.**
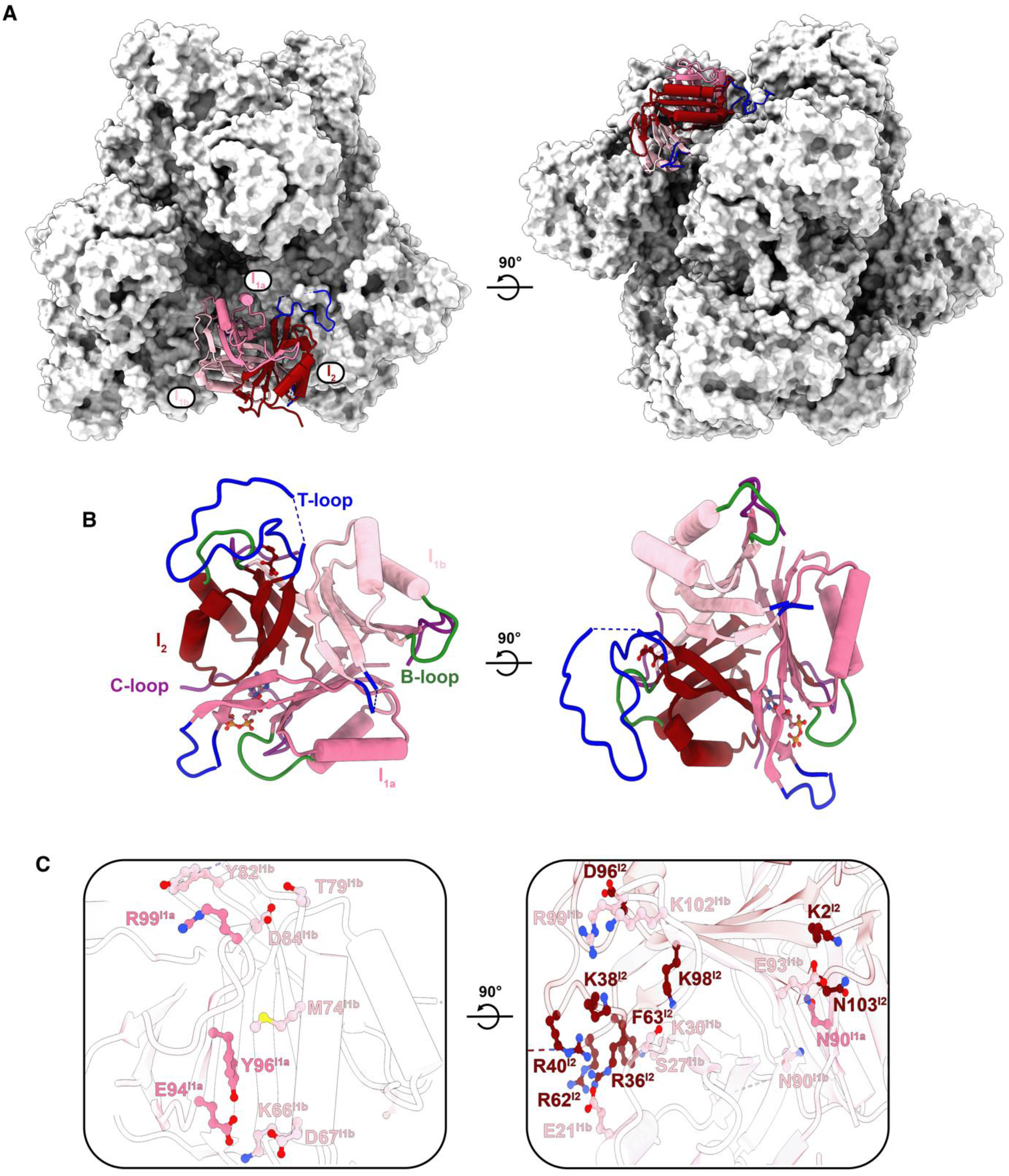
Structural organization and intra-complex interactions within the NifI heterotrimer. **A**. Cartoon representation of a single NifI heterotrimeric complex, composed of NifI_1a_ (pale violet-red), NifI_1b_ (pink), and NifI_2_ (maroon), shown against the grey surface representation of the remaining supercomplex. **B**. Close-up view of the NifI complex illustrating the architecture of NifI_1a_, NifI_1b_, and NifI_2_, with their respective T-, B-, and C-loops colored blue, green, and purple, respectively. **C**. Detailed view of residues mediating intra-complex interactions among the NifI_1a_, NifI_1b_, and NifI_2_ subunits.

### The NifI complexes exhibit differential 2OG and ADP occupancy that reveals functional asymmetry

ATP/ADP and 2OG function as metabolic signals that regulate PII protein interaction with targets^12^. In canonical PII proteins, 2OG binding typically requires co-binding of Mg^2+^ and ATP that results in disassociation from the target protein. Depletion of 2OG has been shown to increase the ATPase activity of PII proteins, converting ATP to ADP that promotes T-loop alteration and association with the target protein^32^. Previous studies with *M. maripaludis* showed that 2OG/ATP promotes the *in vitro* dissociation of the NifI complex from NifDK, establishing the nucleotide/2OG dependent regulation of nitrogen fixation and nitrogenase activity in methanogens^25,26^. As such, ADP-bound NifI complex should be associated with NifDK. However, the binding of ADP and how these ligands engage with the NifI complex when in complex with nitrogenase has not been established. The six NifI complexes in our cryo-EM structure display asymmetric and heterogeneous binding to ADP and/or 2OG (**Fig. 6 & Supplemental Fig. 13**). ADP and/or 2OG interactions are mediated by the B-, C-, and T-loops of the NifI subunits (**Supplemental Fig. 12**). Notably, ADP is found in at least one ligand-binding site in five of the six NifI complexes, consistent with the ADP-bound form driving association with NifDK.

Consistent with the conservation of the ligand binding residues in the primary sequence (**Supplemental Fig. 11**), the architecture of the three ligand binding pockets of the NifI complex is similar to that of GlnK (**Supplemental Fig. 10**). Pocket 1 is formed by the base of the T-loop and B-loop from NifI_1a_ and the C-loop of NifI_2_. Pocket 1 of each NifI complex contains a ligand, with five containing ADP and one with 2OG. Pocket 2 is formed by the base of the T-loop and B-loop from NifI_2_ and the C-loop of NifI_1b_. Pocket 2 displays the most heterogeneity in ligand occupancy, with two NifI complexes containing ADP, two containing 2OG, and two free of any ligands (**Fig. 6**). Pocket 3 is formed by the base of the T-loop and B-loop from NifI_1b_ and the C-loop of NifI_1a_. Pocket 3 is free of ligand in all six NifI complexes (**Fig. 6 and Supplemental Fig. 10**). These results indicate that the ligand affinity differs between the three binding pockets, with pocket three likely having the lowest affinity for ligand. Moreover, the asymmetric binding of ligand by the NifI complexes is a feature of the stable supercomplex. The asymmetry of ligand association may pose the complex for rapid release in response to small changes in 2OG and ATP/ADP levels.

Interestingly, binding of 2OG is tolerated in ligand binding pockets 1 and 2 that still allows stable association of the NifI complex with NifDK. Moreover, 2OG binding by the NifI complexes is independent of ATP. However, previous results with *M. maripaludis* showed that the addition of both ATP and 2OG were required for the *in vitro* disassociation of the NifI complex from NifDK^25^. Similarly, washing resin-bound strep-NifD-NifK with 2OG and ATP removed associated NifI_1_ and NifI_2_ from the complex, resulting in the elution of NifDK that is free of NifI_1_ and NifI_2_ (**Supplemental Fig. 14**). Taken together, these results support the model that ADP binding by the NifI complexes triggers association with NifDK and likely the assembly of the supercomplex. NifI complex binding is reversible with elevated levels of 2OG/ATP releasing the NifI complex from NifDK. The disassociation of the NifI complex from NifDK could lead to disassembly of the supercomplex.

**Figure 6.**
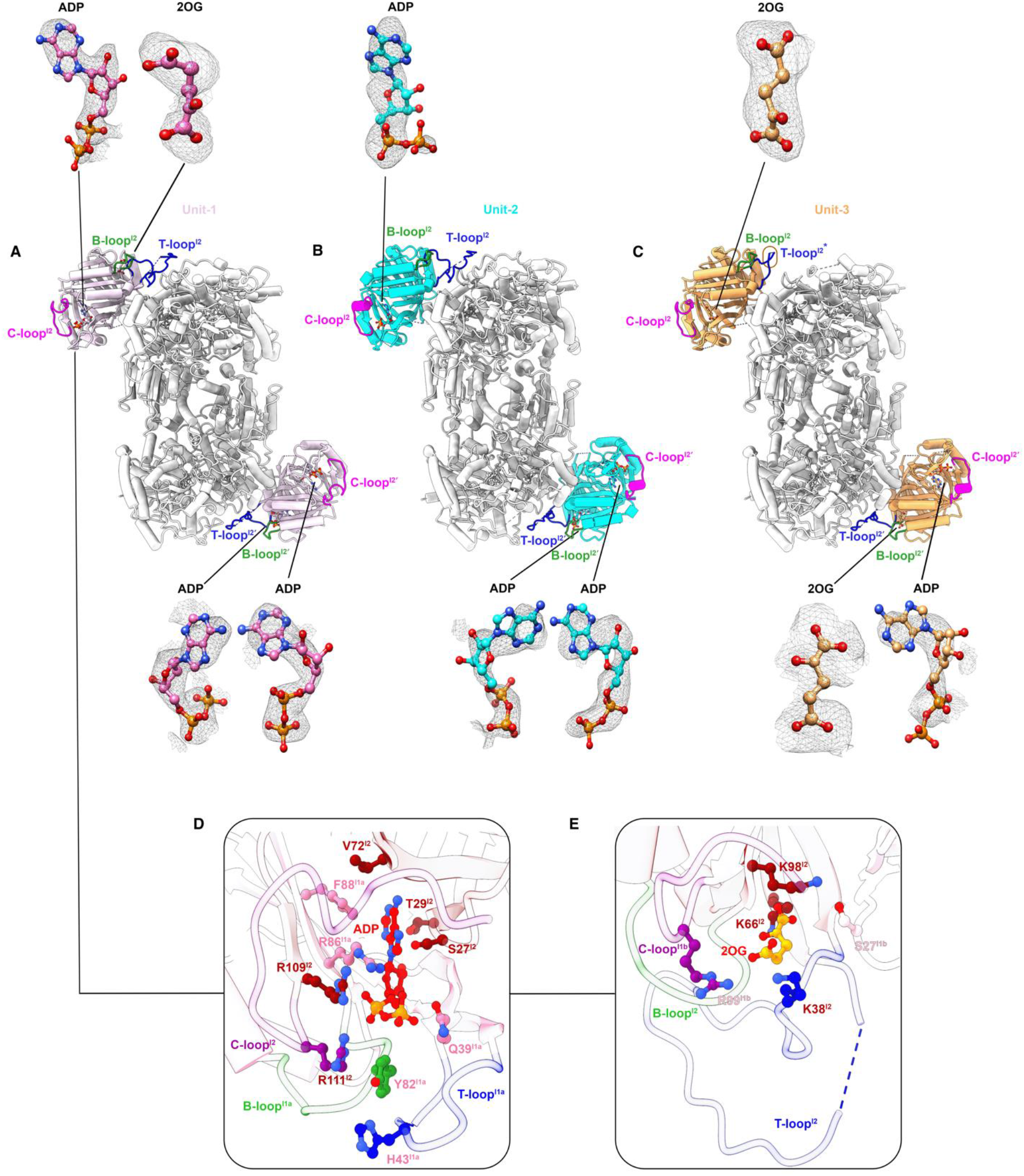
Ligand binding occupancy of the six NifI complexes bound to the three NifDK tetrameric units within the supercomplex. **(A–C)** Structural views of Units 1, 2, and 3 of the supercomplex, respectively, showing the core NifDK subunits in grey and the corresponding NifI complexes in pink, cyan, and sandy brown. For each unit, the T-loop (blue), B-loop (green), and C-loop (magenta) of the NifI_2_ subunit are highlighted. Rectangles indicate the occupancy of ADP or 2-oxoglutarate (2OG) within the NifI complex. The electron density for each of the bound nucleotide/2OG molecules is also depicted. **D**. Detailed view of residues mediating the interaction with ADP in one binding site (left ligand-binding pocket at the top of Unit 1 in panel A) is shown. **E**. Detailed view of residues mediating the interaction between 2OG and the right ligand-binding pocket (right ligand-binding pocket at the top of Unit 1 in panel A) is shown.

### The α-loop insertion of *M. acetivorans* NifD directly interacts with the T-loop of NifI_2_

Although all known and predicted structures of the NifDK proteins are very similar, there is some sequence and structural heterogeneity (**Fig. 7A and Supplemental Fig. 15**). For example, most aerobic diazotrophic bacteria have an N-terminal extension in their NifK (β) subunit that is hypothesized to stabilize the heterotetramer^33^. In contrast, not only do diazotrophic bacteria and archaea that are anaerobes lack the NifK extension, but many instead have a ∼ 50 amino acid insertion within NifD (α-subunit) called the α-loop insertion^33^ (**Fig. 7A**). For example, the crystal structure of NifDK from *Clostridium pasteurianum* shows the α-loop insertion adjacent to the predicted binding site of NifH, indicating that the α-loop is a specificity determinant for interaction with NifH^34^. All methanogens known to or predicted to fix nitrogen possess a *nif* operon that contains both NifI_1_ and NifI_2_, indicating that NifI-dependent control of nitrogenase activity is a universal feature of nitrogen fixation in methanogens^16^. However, deeply rooted methanogens, such as *M. maripaludis*, lack the α-loop insertion within NifD while later evolving Methanosarcinales, including *M. acetivorans*, have the NifD α-loop insertion (**Supplemental Fig. 15**)^33^. However, *C. pasteurianum* lacks NifI proteins.

**Figure 7.**
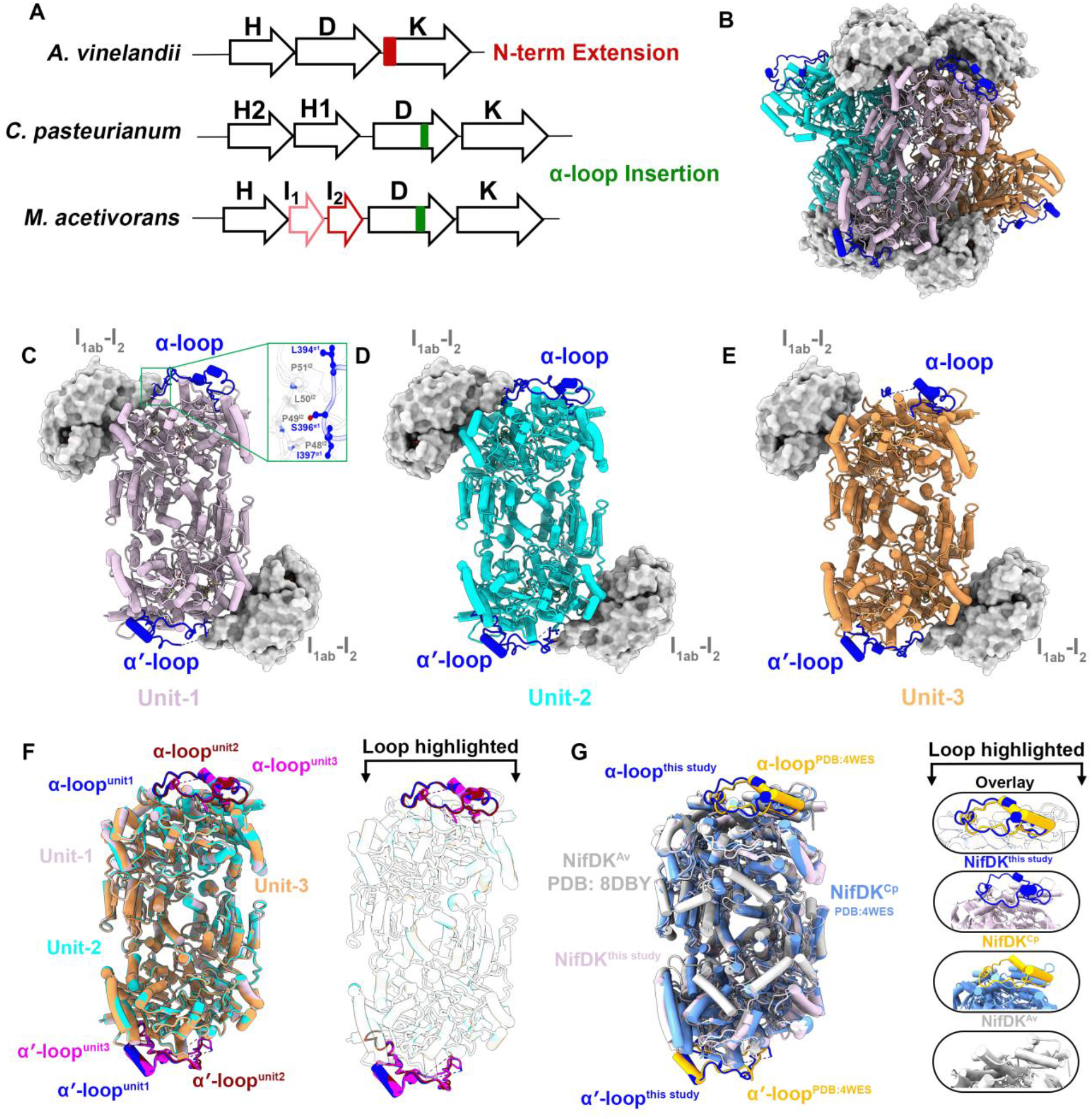
α-loop features in NifDK. **A**. Schematic comparison of NifDK subunit architecture in *Azotobacter vinelandii* (Av), *Clostridium pasteurianum* (Cp), and *Methanosarcina acetivorans* (Ma), highlighting the N-terminal extension in *Av* (red) and α-loop insertion in *Cp* and *Ma* (green). **B**. Overall structure of the three NifDK tetrameric units (Unit 1, pink; Unit 2, cyan; Unit 3, orange) with α- and α′-loops (blue) mapped. The NifI complexes are surface-rendered (grey). **C–E**. Detailed structures of NifDK and NifI complexes in Unit 1, Unit 2, and Unit 3, respectively, with α- and α′-loops highlighted. Inset in panel C showcases residues in the α-loop of Unit 1 and NifI_2_ that promote the interaction. **F**. Superposition of the three NifDK tetrameric units illustrating variability in α- and α′-loop conformations. **G**. Structural overlay of the α- and α′-loops in this study (blue) with those from *C. pasteurianum* (PDB: 8DBY) and *A. vinelandii* (PDB: 4WES), showing loop conformational diversity.

In our supercomplex structure, not only does the NifI complex binding site overlap with the putative NifH binding site (**Fig. 2B**), but the T-loop of NifI_2_ also extends out and associates directly with the NifD α-loop insertion in five of the six NifI complexes (**Figs. 6, 7 & Supplemental Fig. 16**). Thus, the α-loop is a key specificity determinant in the association of the NifI complex with NifDK in *M. acetivorans*. This result is consistent with substantial differences in the NifI_2_ T-loop sequences between *M. acetivorans* and *M. maripaludis* (**Supplemental Fig. 15**). Indeed, the structure of the PII-NifDK supercomplex from *Methanocaldococcus infernus* reveals distinctly different NifI complex binding to NifDK, and supercomplex architecture, indicating supercomplex formation is a universal feature in methanogens, but supercomplex is different between methanogens that contain or lack the α-loop insertion. The T-loop of *M. acetivorans* NifI_2_ is uniquely suited to interact with the α-loop insertion, and this interaction is likely depending on ligand binding by the NifI complex. Specifically, binding of ADP is likely critical for facilitating the NifI_2_ T-loop in an extended confirmation to allow the association with the α-loop of NifDK. Notably, the only NifI complex with an unstructured NifI_2_ T-loop and a NifDK α-loop is the NifI complex that lacks any ADP and contains only 2OG in ligand binding pocket 2 (**Fig. 7E**). Moreover, the loss of the T-loop/α-loop interaction decreases the buried surface area between the NifI complex and NifDK in this unit by one-half (**Supplemental Fig. 8C**), indicating that this interaction is substantial but not essential for the stable association of the NifI complex to NifDK, at least in the supercomplex. The shorter NifI_1a_ T-loop also has a structured confirmation. However, the T-loop of NifI_1b_ in all NifI complexes is unstructured, indicating that this T-loop is not involved in association of the NifI complex with NifDK in the supercomplex. These results reveal that the T-loop of NifI_2_ and the α-loop of NifD are critical and unique factors involved in the association of the NifI complex with NifDK in *M. acetivorans*.

### Steps in the supercomplex assembly/disassembly process

In addition to the supercomplex, our cryo-EM datasets of the natively isolated Strep-NifD sample possessed three additional smaller 2D classes (**Supplemental Figs. 2, 3 & 4; Supplemental Table 2**). These classes gave rise to three states of NifDK structures (**Fig. 8)**: D*KKD* (State-1; **Supplemental Figure 17; Supplemental Table 2**), DKK (State-2; **Supplemental Figure 18; Supplemental Table 2**), and a PII-bound DKKD* complex (State-3; **Supplemental Figure 19; Supplemental Table 2**). In all these structures, the two NifK subunits are ordered and the density for the P-cluster is intact. The overall structural details of NifK and the P-cluster are similar to those in the supercomplex structure (State-4, **Fig. 8**). However, density for either one or both NifD subunits are poor (denoted by the * nomenclature). In State-2 and State-3, the M-cluster is clearly visible in the ordered NifD subunit, and the structural details are similar to those of the supercomplex. In addition, the PII complex in State-3 is bound to three ADP molecules (**Supplemental Figure 19**). These data are consistent with nucleotide binding and hydrolysis states regulating PII binding to the NifDK tetramer and formation/dissociation of the supercomplex.

**Figure 8.**
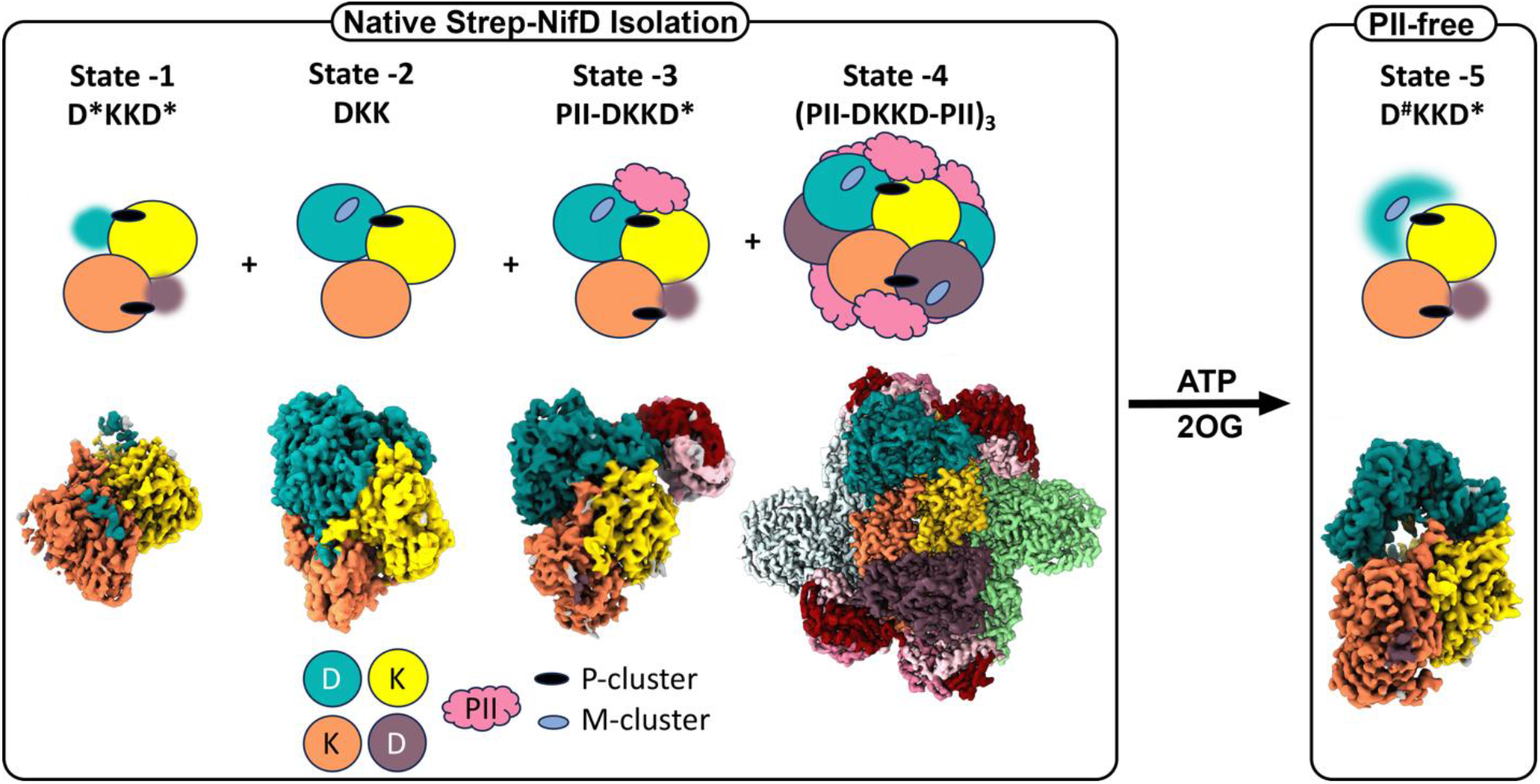
Structural snapshots of the assembly/disassembly complexes. Cartoon representation of the four states captured in this Cryo-EM study and their corresponding structures are shown. D* represents presence of partial density for the NifD subunit. When PII is removed upon addition of ATP and 2OG, only State-5 is observed in Cryo-EM analysis. D^#^ represents an arch-like conformation for one NifD subunit and D* denotes partial density for the other subunit.

Mass photometry of the Strep-NifD sample reveal that both the supercomplex and tetrameric forms are present (**Supplementary Figs. 20A & B**). The broad nature of the peak (230±105 kDa) in Supplementary Fig. 20B suggests that there are multiple states of NifDK, including those bound to PII complex. These data are consistent with the multiple states observed in the cryo-EM data. Addition of 2OG and ATP to the sample leads to a reduction in the tetrameric population with a subsequent accumulation of a broader, but smaller species (117±21 kDa; **Supplementary Fig. 20C**). Complete removal of NifI from the sample through on column treatment of the Strep-NifD pull down sample with 2OG and ATP (**Supplemental Fig. 14**) results in the detection of primarily the NifDK tetramer (∼239±35 kDa), two smaller species (169±24 kDa and 101±22 kDa), and a small population of a ∼437±71 kDa species, as seen in mass photometry analysis (**Supplementary Fig. 20D**). These data suggest that the PII complex might increase the structural rigidity of the tetramer towards promoting supercomplex formation. To further test this model, we assessed the NifI-free Strep-NifD sample using cryo-EM. Here, only one 2D class was observed (**Supplementary Fig. 21**) and this resulted in an additional D^#^KKD^*^ structure (State-5; **Fig. 8 & Supplemental Fig. 22**). No classes corresponding to the supercomplex (State-4), or States 1-3 were observed. Density for one NifD in this complex is poor (D*) and the other NifD is ordered but forms an arch-like conformation (D^#^).

## DISCUSSION

Our cryo-EM structures reveal a previously uncharacterized mechanism of nitrogenase regulation in methanogens, involving the formation of a higher-order complex through direct interactions between NifDK catalytic subunits. The non-canonical PII proteins NifI_1_ and NifI_2_ assemble into a heterotrimeric complex that binds each half of the NifDK heterotetramer, resulting in (1) inhibition of nitrogenase activity by blocking electron transfer from NifH, and (2) formation of an inactive supercomplex comprising three NifDK units.

Given that all known and predicted diazotrophic methanogens encode NifI_1_ and NifI_2_, this supercomplex likely represents a conserved regulatory feature in these organisms. Indeed, similar-sized complexes were observed by size-exclusion chromatography after purification of NifI_1,2_-NifDK from *M. maripaludis*^25^. While no analogous structures have been observed in bacteria, the presence of NifI homologs in several diazotrophic bacterial species suggests that similar regulatory mechanisms may exist outside of methanogens^10,35^. Notably, NifI proteins are restricted to anaerobes, raising the question of why anaerobic organisms - including all methanogens - utilize NifI-mediated regulation, whereas aerobes rely on alternative systems to control nitrogenase activity.

Methanogens operate near the thermodynamic limits of life, deriving minimal energy from methanogenesis compared to aerobic respiration<u>^17^</u>. Consequently, energy-efficient regulation of nitrogen fixation is essential. Direct inhibition of NifDK by PII complex binding in response to fixed nitrogen availability is likely more rapid and energetically favorable than indirect mechanisms. Supporting this, ammonia addition leads to a swift loss of nitrogenase activity in methanogens^<u>20,</u>22^. In contrast, many aerobic bacteria employ the DraTG system, which inactivates nitrogenase via ADP-ribosylation of NifH - a process that requires additional enzymes and substrates and is likely slower and less efficient^11,13^.

The formation of the supercomplex appears to stabilize the inactive state of NifDK beyond simply preventing NifH binding. Two lines of evidence support this: (1) the NifI complex bridges two NifDK heterotetramers, increasing the interaction surface and enhancing complex stability, as evidenced by successful purification of the supercomplex,; and (2) recent cryo-EM studies of bacterial nitrogenase reveal conformational changes in NifDK during catalysis^36^, which are likely restricted in the PII-NifDK supercomplex. This structural constraint may further prevent catalysis and limit substrate access to the active site. Importantly, PII complex binding is reversible, as shown by dissociation from resin-bound NifDK and increased activity in the presence of 2OG and ATP.

Unexpectedly, in the native state, we observed asymmetrical binding of ADP and 2OG within the NifI complexes. In canonical PII proteins, ADP binding typically promotes target association, while MgATP-2OG binding typically promotes dissociation^12^. Here, 2OG binds in the absence of ATP, indicating that partially 2OG occupancy is insufficient to trigger dissociation of the NifI complex. Ligand occupancy was detected in binding pockets 1 and 2 - each partially formed by NifI_2 -_ while pocket 3, formed between NifI_1a_ and NifI_1b_, remained unoccupied. Despite this asymmetry, the T-loops across the six NifI complexes are largely symmetric, with all six NifI_1b_ T-loops disordered/unstructured, all six Nif_1a_ T-loops structured, and five of the six longer NifI_2_ T-loops structured (**Fig. 6, Supplemental Figs. 10 & 12**). The lone exception is the disordered/unstructured NifI_2_ T-loop in the top half of Unit 3, which is the only NifI complex to have only 2OG bound (**Fig. 7)**. This asymmetry suggests a stable feature of the supercomplex, potentially allowing it to respond more sensitively to fluctuations in ATP/ADP and 2OG levels than a uniformly ADP-bound complex.

The T-loop of canonical PII proteins plays a central role in both ligand binding and target interaction. Consistent with this, deletion of the T-loop region in either NifI_1_ or NifI_2_ from *M. maripaludis* disrupts co-purification with NifDK - completely abolishing it with NifI_1_-ΔT and significantly reducing it with NifI_2_-ΔT^25^. The *M. acetivorans* PII-NifDK supercomplex supports these findings. While the NifI_1a_ and NifI_1b_ T-loops do not directly contact NifDK, the NifI_1a_ T-loop is essential for ligand binding at pocket 1, which is consistently occupied by ADP or 2OG across all six NifI complexes (**Supplemental Fig. 10**). This suggests that ligand occupancy at pocket 1 is required for NifI complex association with NifDK, and without the NifI -T-loop ligand cannot be bound. *M. maripaludis* NifI_2_-ΔT retains some binding to NifDK, consistent with the *M. acetivorans* supercomplex structure that reveals NifI_2_ contributes most of the interaction surface with NifDK, with approximately half of it located outside the T-loop (**Supplemental Fig. 8**).

Importantly, the NifI_2_ T-loop directly interacts with the α-loop insertion in NifD (**Fig. 7**), identifying the α-loop as a key specificity determinant for NifI complex interaction with NifDK. This insertion is absent in *M. maripaludis* and other Methanococci but present in all Methanosarcinales and in *Clostridium pasteurianum*, although the latter lacks NifI_1_ and NifI_2_ and shows significant sequence divergence in the α-loop (**Supplemental Fig. 15**). Despite sequence differences, the α-loops of *M. acetivorans* NifD and *C. pasteurianum* NifD are structurally conserved (**Fig. 7F**), suggesting a conserved structural role. The NifI_2_ T-loop to NifD α-loop interaction appears to be a defining feature of nitrogenase regulation in Methanosarcinales. In contrast, Methanococci rely on alternative NifD regions for NifI_2_ T-loop interaction^37^. The acquisition of the α-loop insertion in Methanosarcinales may have evolved to enhance specificity of NifDK for NifI and NifH, particularly given their unique capacity to express all three nitrogenase isoforms - Mo-, V-, and Fe-nitrogenases^18^. We previously demonstrated that *M. acetivorans* expresses all three simultaneously^18^, and the α-loop likely ensures selective interaction of NifH and NifI with NifDK, excluding VnfDGK and AnfDGK. Supporting this, the Vnf and Anf operons in *M. acetivorans* encode distinct PII proteins with divergent T-loops (**Supplemental Fig. 11**), despite each using the same Fe-protein^15^. Similarly, given the absence of NifI proteins, the α-loop in *Cp*NifD may serve to restrict NifH interaction to NifDK. Thus, the supercomplex structure highlights the coevolution of nitrogenase (e.g. α-loop insertion) with factors (NifI complex) that control activity.

Finally, this study uncovers a previously unrecognized mechanism of nitrogenase regulation in methanogens, characterized by the formation of a stable, inactive supercomplex mediated by non-canonical PII proteins NifI_1_ and NifI_2_. The direct inhibition of NifDK, coupled with structural stabilization through higher-order assembly, represents an energy-efficient strategy uniquely suited to the metabolic constraints of anaerobic life. The discovery of the α-loop insertion in NifD as a specificity determinant for NifI_2_ interaction further reveals a molecular basis for selective nitrogenase regulation in *Methanosarcinales*, particularly in organisms expressing multiple nitrogenase isoforms. These findings expand our understanding of nitrogenase control, highlight evolutionary adaptations in anaerobic archaea, and suggest broader implications for nitrogen fixation strategies across diverse microbial lineages.

## Acknowledgments

This work was supported by grants from the Department of Energy, Office of Basic Energy Sciences, DE-SC0020965 (EA) for structure determination, DE-SC0019226 (DJL) for strain construction, protein purification, and activity analyses, DE-SC0018143 (BB) for mass spectrometry. We thank Drs. Summer Brock, Katherine Basore, and Bradley Readnour at the Washington University Center for Cellular Imaging and Drs. Liguo Wang, Guobin Hu, and Jake Kaminsky at the Laboratory for BioMolecular Structure (LBMS) at Brookhaven National Labs for Cryo-EM data collection. LBMS is supported by the Department of Energy, Office of Biological and Environmental Research (KP1607011). Funding for the Montana State Mass Spectrometry Facility used in this publication was made possible in part by the MJ Murdock Charitable Trust, MSU office of the VPRED, and the National Institute of General Medical Sciences of the National Institutes of Health under Award Numbers P20GM103474 and S10OD28650. We also acknowledge generous financial support from the Doisy Research Fund of the Edward A. Doisy Department of Biochemistry and Molecular Biology at Saint Louis University School of Medicine.

## Author contributions

Conceptualization: EA and DJL

Methodology: RK, TMD, AD, MC, MTK, and BB

Investigation: RK, TMD, AD, MC, and MTK

Visualization: RK, TMD, EA, and DJL

Funding acquisition: EA, DJL, BB

Project administration: EA and DJL

Supervision: EA, DJL, and BB

Writing – original draft: DJL, EA, and RK

Writing – review & editing: all authors

## Data and materials availability

The single particle cryo-EM maps and models have been deposited into the PDB and EMDB with the following codes: State-1 (D*KKD*) PDB:11UY & EMD-76071. State-2 (DKKD*) PDB 11SX & EMD-76025. State-3 (PII-DKKD*) PDB 11MY & EMD-75852. State-4 (Supercomplex (C1 symmetry), (PII-DKKD-PII)_3_ PDB: 9P1X & EMD-71144. State-4 (Supercomplex (C3 symmetry), (PII-DKKD-PII)_3_ PDB: 12AH & EMD-70378. State-5 (D^#^KKD*) PDB 11ZK & EMD-76217.

A Source Data File is also available and contains the raw data used for activity measurements and the mass-spectra analysis of the bands from the pull-down experiment.

## Supplemental Materials

Supplemental Figs. 1 to 22

Supplemental Tables 1 to 5

## Methods

### Cell Growth

*M. acetivorans* strains were grown in anoxic high-salt (HS) medium at 35º C as previously described^19^. HS medium was prepared inside an anaerobic chamber (Coy Laboratories) containing 75% N_2_, 20% CO_2_, and 5% H_2_. The medium was supplemented with 2 µg/mL puromycin when required. Growth comparison experiments were performed in Balch tubes containing 10 ml of HS medium containing 125 mM methanol, 1 mM sulfide and 3 mM cysteine. Fixed nitrogen (18 mM NH_4_Cl) was added from anaerobic sterile stock solutions prior to inoculation where indicated. The optical density was measured at 600 nm using a spectrophotometer. For purification of Strep-NifD, *M. acetivorans* cells were grown in 1.5 L HS medium lacking NH_4_Cl with 125 mM methanol and 0.025% Na_2_S. Cells were grown to late log phase (OD_600_ = ∼0.5), harvested anaerobically by centrifugation, and the pellet was stored at -80°C in an anoxic Hungate tube. *E. coli* strains were grown and maintained with solid or liquid LB medium with the appropriate antibiotic. For recombinant Strep-NifH purification, *E. coli* SufFeScient cells^38^ containing pDL906 were grown in four liters of LB medium with 100µg/mL kanamycin, 1 mM L-cysteine, and 1 mM ferric citrate, shaking at 240 rpm at 35º C. At an OD_600_ of 0.5-0.6, expression was induced with 0.2 mM IPTG and supplemented with 0.5 M D-sorbitol and set to shake at 150 rpm at 16º C. After 16 hours, cultures were transferred to sterile GL45-closure reagent bottles with 1.7 mM sodium dithionite, sealed tightly, and incubated at 16º C for 3 hours. These cells were then harvested anaerobically by centrifugation, and the pellets stored at -80°C in anoxic containers.

### Strain Construction

Plasmids, primers, and strains used in this study are listed in **Supplemental Tables 3 & 4**. *M. acetivorans* strain DJL70 that expresses Strep-NifD was generated using the CRISPR-Cas9 system^39^ with a few modifications. All primers and gBlocks were designed using Geneious Prime software and purchased from IDT. The gRNA-*nifI*_*2*_ was designed to target Cas9 to cut within *nifI*_*2*_ (MA3897) upstream of *nifD* (MA3898). Homology regions upstream and downstream of *nifD* were obtained as synthetic gBlocks and contained sequence to add a twin-Strep-tag with linker, identical to as described for Mcr^40^, to the N-terminus of NifD. A single nucleotide change (C to A) in the PAM sequence was added to prevent further Cas9 targeting of DNA by gRNA-*nifI*_*2*_. The C to A change does not alter the amino acid sequence of NifI_2_. The homology repair templates and gRNA-*nifI*_*2*_ were assembled into *Asc*I-digested pDL238 using Gibson assembly Ultra Kits (Telesis Bio) followed by transformation into *E. coli* WM4489 competent cells. Transformants were screened by PCR using primers P7.3 and P6.53 to confirm insert assembly into pDL238. The resultant plasmid pDL705 was retrofitted with pAMG40 using Gateway BP Clonase II enzyme mix (Invitrogen) followed by transformation into WM4489 cells. Transformants were screened by PCR using the primer set P6.64 /P6.65 and the resultant plasmid was designated pDL707.

*M. acetivorans* WWM73 cells were transformed with pDL707. Transformants were selected on anaerobic HS agar plates containing 125 mM methanol and 2 µg/mL puromycin as previously described<u>^41^</u>. Colonies were screened by PCR using primers P7.20 and P7.18 that anneal to the Strep-tag and the 3’-end of *nifD*. Cells from a positive colony were passed through HS medium lacking puromycin then plated on HS agar plates containing 125 mM methanol and 50 µg/mL 8-aza-2,6-diaminopurine sulfate (8-ADP) as described^39^ to select for the loss of pDL707. Colonies were assessed for the loss of pDL707 by PCR using primers P7.45 and P7.46. The resultant plasmid-free and puromycin-sensitive strain was designated strain DJL70. PCR amplification of *nifI*_*2*_-*strep-nifD* from the chromosome of strain DJL70 using primers P7.14 and P7.18, followed by sequencing of the PCR product confirmed the addition of the twin-Strep-tag to NifD.

An *E. coli* strain capable of expressing Strep-tagged NifH (Strep-NifH) was generated by PCR amplifying *nifH* from *M. acetivorans* C2A genomic DNA using Q5 polymerase and primers P8.67 and P8.43 to add restriction sites for *Nco*I and *Xho*I as well as a N-terminal strep tag. The PCR product was digested with *Nco*I and *Xho*I according to manufacturers’ instructions (NEB) and ligated into *Nco*I/*Xho*I-digested pET28a using T4 DNA ligase and incubated for 16 hours at 16°C. The ligation reaction was used to transform *E. coli* DH5α cells and selection done using LB agar plates containing 100 μg/mL kanamycin. Colony PCR and sequencing confirmed *strep-NifH* insertion into pET28a, resulting in plasmid pDL906. Plasmid pDL906 was used to transform *E. coli* “SufFeScient” cells and transformants selected by plating on LB agar plates containing 100 μg/mL kanamycin. *E. coli* “SufFeScient” (pDL906) was used for recombinant production and purification of Strep-NifH.

### Protein Purification

All purification steps were performed under anoxic conditions unless otherwise stated. For purification of Strep-NifD, a *M. acetivorans* strain DJL70 cell pellet was resuspended in resuspension buffer (50 mM Tris, 150 mM NaCl pH = 7.2 with 1 mM each PMSF and benzamidine HCl, plus 1.7 mM sodium dithionite). Cells were lysed by sonication on ice with a Qsonica Q55 using a CL-188 probe. Five pulses of sonication (10 seconds each) at an amplitude of 30% with at least one minute on ice between pulses) was followed by anaerobic centrifugation for 10 minutes at 16,000 x *g*, and the supernatant was passed through a 0.45 µm filter. Cleared lysate was loaded onto IBA Strep-Tactin Superflow resin 2 times. The column (0.5 mL) was washed 5 times with Buffer W (50 mM Tris, 150 mM NaCl pH = 7.2 with 1.7 mM sodium dithionite) and protein eluted with Buffer E (Buffer W with 2.5 mM desthiobiotin). Elutions were concentrated using Nanosep 10K Omega columns according to manufacturers’ instructions (Pall Corp.).

For purification of NifI-free NifDK complex, the cleared lysates were applied on to the Strep-Tactin column as described above. Subsequently, after loading, the column was subjected to sequential washes to dissociate NifI. Following initial washing (5 × 1 mL Buffer W), the column was washed with Buffer W supplemented with 10 mM 2-oxoglutarate (5 × 0.5 mL), followed by Buffer W containing 10 mM 2-oxoglutarate, 10 mM MgCl_2_, and 10 mM ATP (5 × 0.5 mL). The column was then equilibrated with Buffer W (5 washes) prior to elution with Buffer E. Eluted fractions were flash-frozen without further concentration.

For purification of Strep-NifH, a SufFeScient (pDL906) cell pellet was resuspended in Resuspension Buffer plus 10 μg/mL DNase I (Sigma) before lysis by three passages through a French pressure cell at > 1100 psi. Lysate was centrifuged at 39,000 x *g* and 4 °C for 10 min, then the supernatant was passed through a 0.45 µm filter. Strep-Tactin column loading, washing with Buffer W, and elution with Buffer E were performed as described above for Strep-NifD.

Strep-NifD and Strep-NifH proteins were quantitated using a Qubit 2.0 fluorometer (Invitrogen). Protein purity was analyzed by SDS-PAGE using a 15% gel (**Supplemental Fig. 1**). A Broad Range (10–250 kDa) color prestained protein standard (New England Biolabs) was used to approximate the molecular weight of each protein. SDS-PAGE gels were stained with Coomassie Brilliant Blue solution, then destained prior to imaging. The protein bands were excised and sent for trypsin digestion followed by mass spectrometry analysis by the IDeA National Resource for Quantitative Proteomics facility at University of Arkansas for Medical Sciences (Source data file).

### Nitrogenase activity assay

Nitrogenase activity was assayed by measuring reduction of acetylene to ethylene using methods similar to those described previously^25^. The assay mixtures contained 25 mM HEPES, pH = 7.5, with 5 mM ATP, 12.5 mM MgCl_2_, 25 mM creatine phosphate, 100 U/mL creatine phosphokinase (Sigma), 20 mM sodium dithionite in reaction volumes of 1 mL. Where indicated, these mixtures were supplemented with 10 mM 2-oxoglutarate. All reactions contained 50 μg purified Strep-NifD and were initiated by the addition of 50 μg purified Strep-NifH. Reaction vials were 10 mL serum vials (Kimble) with a total internal volume measured at 13 mL. These were closed with butyl rubber stoppers and sealed with 20 mm crimps before purging with argon for 5 minutes. 2 mL of acetylene was added by syringe to each vial, then 800 μL assay mix was dispensed by needle to each, and reactions initiated by adding 200 μL Strep-NifH solution by syringe. Ethylene production was monitored by periodic sampling of 200 μL headspace which was injected onto a GC-2030 gas chromatograph (Shimadzu) with a CP-PoraPLOT Q capillary column (Varian, #CP7551). Ethylene standards were used to quantify results.

### Western blot

The production of Strep-NifD in *M. acetivorans* strain DJL70 was assessed by the detection of Strep-NifD using a custom polyclonal NifD-specific antibody and a monoclonal Strep-tag-specific antibody (Qiagen, #34850) as previously described^19^.

### Cryo-EM Sample Preparation and Data-Collection

Cryo-EM samples were prepared following a protocol previously developed for the nitrogenase-like enzyme DPOR^42^. The eluted NifD protein sample (0.7 mg/ml), which co-purifies with NifK, NifI_1_, and NifI_2_, was diluted to 0.4 mg/ml using Buffer W in a sealed glass vial inside an anaerobic chamber. Quantifoil R2/2 holey carbon grids (Q350CR2, Electron Microscopy Sciences) were negatively glow discharged for 60 seconds using a Gloqube (Quorum). Immediately following grid cleaning, 3 µL of the protein sample was applied to each grid using a Hamilton gas-tight syringe to minimize air exposure. Grid preparation was performed using a Vitrobot Mark IV (Thermo Fisher Scientific) with the chamber maintained at 4°C and 100% humidity. Without additional incubation, grids were blotted for 2.5 seconds (blot force 2) and plunge-frozen into liquid ethane cooled by liquid nitrogen. The total exposure time of each sample to ambient air during transfer and freezing was less than 30 seconds. Grids were stored in liquid nitrogen until data collection. This preparation strategy consistently yielded thin, uniform vitreous ice with no evidence of air-water interface artifacts. The NifI-free NifDK protein sample (final concentration 0.4 mg/ml) was prepared and handled similarly.

Grids were clipped and loaded into either a 200 kV Thermo Fisher Scientific Glacios or a 300 kV Thermo Fisher Scientific Krios transmission electron microscope, as specified for each dataset in Supplementary Tables 3 and 4. Data were acquired using EPU software (Thermo Fisher Scientific) with aberration-free image shift (AFIS) and recorded on a Falcon IV direct electron detector (4k × 4k). Imaging parameters, including pixel size, defocus range, spherical aberration, and total electron dose, varied between datasets and are summarized in Supplementary Tables 3 and 4. All datasets exhibited good particle distribution and orientation diversity, as assessed by 2D classification.

### Data processing

#### State-4

Single-particle cryo-EM data were processed using cryoSPARC v5.0.2. For the NifDK dataset, 3,826 raw movies were motion-corrected using patch motion correction, followed by contrast transfer function (CTF) estimation via patch CTF estimation (**Supplemental Fig. 2**). Initial particle picking was performed using the blob picker, and particles were extracted with a box size of 400 pixels (binning factor 4). This yielded an initial stack of 1,798,035 particles. Multiple rounds of 2D classification were conducted to remove false positives and damaged particles, resulting in a cleaned subset of 899,239 particles. These particles were used to generate *ab-initio* 3D reconstructions. The particle stack corresponding to the heterotrimeric complex (124,698 particles) underwent multiple rounds of 2D classification to remove poorly aligned particles and improve homogeneity. An initial 3D reconstruction was generated using *ab-initio* modeling in cryoSPARC, followed by non-uniform refinement to enhance resolution and account for structural variability. Subsequent 3D classification resolved conformational heterogeneity, particularly differences in particle populations based on the presence or absence of associated subunits. Among the resulting classes, one, comprising the largest number of particles, yielded the highest-quality 3D reconstruction and clearly resolved all three components of the heterotrimeric complex. This class was further refined using homogeneous refinement followed by non-uniform refinement, with particles re-extracted at full resolution (binning factor of 1). The final map, generated from a subset of 74,083 particles, reached a resolution of 3.16 Å at a Fourier shell correlation (FSC) threshold of 0.143 and was further sharpened using DeepEMhancer (**Supplemental Table 1**).

### Initial preprocessing (states 1 and 3)

For the dataset used to resolve D*KKD* (*state 1*) and DKKD*-PII (*state 3*), a total of 13,077 raw movies were motion-corrected using patch motion correction, and contrast transfer function (CTF) parameters were estimated using patch CTF estimation (Supplemental Fig. 3). Initial particle picking with a blob picker and extraction (box size 300 pixels, binning factor 4) yielded 4,718,455 particles. Multiple rounds of 2D classification removed false positives and damaged particles, resulting in a cleaned stack of 4,640,379 particles.

#### State-1

From the cleaned stack, particles corresponding to D*KKD* were selected and processed using the HR-HAIR workflow optimized for smaller volumes (200 classes, initial uncertainty factor 1, circular mask diameter 150 pixels, 20 final iterations, 80 online EM iterations, batch size 400 particles per class). Classes displaying well-defined features were selected for ab-initio 3D reconstruction. One resulting class (217,846 particles) displayed high-resolution features and was re-extracted at full resolution without binning for homogeneous reconstruction with gold-standard splitting and local refinement (rotation search 2°, shift search 1 Å, recentering at each iteration, initial low-pass filter 6 Å). The final map, generated from 217,545 particles, reached an overall resolution of 3.12 Å (FSC = 0.143) and was further sharpened using DeepEMhancer (Supplemental Figure 3 & Table 4).

#### State-2

A second particle population corresponding to DKK bound to NifI_1ab_ and NifI_2_ was identified in the original dataset used for supercomplex reconstruction (334,817 particles). These particles were processed using the HR-HAIR workflow with the same parameters as above. Ab-initio reconstruction was carried out with three 3D classes (initial resolution 6 Å, maximum resolution 4 Å, Fourier radius step 0.005, initial minibatch size 300, final minibatch size 1000, real-space centering disabled). One class (135,383 particles) displayed well-defined structural features and was re-extracted at full resolution for homogeneous reconstruction with gold-standard splitting and local refinement (rotation search 2°, shift search 1 Å, recentering at each iteration, initial low-pass filter 6 Å). The final map, generated from 135,383 particles, reached an overall resolution of 3.33 Å (FSC = 0.143) and was further sharpened using DeepEMhancer (Supplemental Figure 4 & Table 4).

#### State-3

From the same cleaned particle stack used for state 1, particles corresponding to PII-DKKD* (404,214) were selected for further processing. Subsequent 2D classification and ab-initio reconstruction were carried out using the HR-HAIR workflow optimized for smaller volumes (300 classes, initial uncertainty factor 1, circular mask diameter 200 pixels, 20 final iterations, 80 online EM iterations, batch size 400 particles per class, maximum reconstruction resolution 3 Å). One class (175,275 particles) displayed high-resolution features and was re-extracted at full resolution for homogeneous reconstruction with gold-standard splitting and local refinement (rotation search 2°, shift search 1 Å, recentering at each iteration, initial low-pass filter 6 Å). The final map, generated from 175,013 particles, reached an overall resolution of 3.03 Å (FSC = 0.143; Supplemental Figure 3 & Table 4).

#### State-5

A total of 18,000 raw movies were motion-corrected using patch motion correction, and contrast transfer function (CTF) parameters were estimated using patch CTF estimation (**Supplemental Fig. 21**). Initial particle picking with a blob picker and extraction (box size 400 pixels, binning factor 4) yielded 5,698,344 particles. Multiple rounds of 2D classification removed false positives and damaged particles, resulting in a cleaned stack of 4,535,974 particles. Subsequent 2D classification was performed using the HR-HAIR workflow optimized for smaller volumes (200 classes, initial uncertainty factor 1, circular mask diameter 120 pixels, 20 final iterations, 80 online EM iterations, batch size 400 particles per class)<u>^43^</u>. Classes displaying well-defined high-resolution features were selected for *ab-initio* 3D reconstruction. A first round of HR-HAIR *ab-initio* reconstruction was carried out with three classes (initial resolution 6 Å, maximum resolution 4 Å, Fourier radius step 0.005, initial minibatch size 300, final minibatch size 1000, real-space centering disabled). One class (1,635,266 particles) showed clear high-resolution features and was re-extracted at full resolution without binning for a second round of HR-HAIR 2D classification. In the second round, 2D classification was performed using the same parameters, yielding refined classes (1,457,524 particles) that were selected for a second *ab-initio* reconstruction. Using identical reconstruction settings, one class (591,856 particles) displayed high-resolution features and was subjected to homogeneous reconstruction with gold-standard splitting and local refinement (rotation search 2°, shift search 1 Å, recentering at each iteration, initial low-pass filter 6 Å). The final map, generated from 591,856 particles, reached an overall resolution of 3.01 Å and was further improved by map sharpening using DeepEMhancer (FSC = 0.143; **Supplemental Table 2**). All particle populations observed in the initial dataset, including the higher-order NifDK–NifI heterotrimeric supercomplex and DKK species, were also present in the expanded dataset.

### Mass Photometry

All measurements were carried out on a TwoMP instrument (Refeyn Ltd.). Glass coverslips (No. 1.5H thickness, 24 × 50 mm, VWR) and holey silicone gaskets (Grace Bio-Lab’s reusable CultureWell gaskets, Sigma) were cleaned by sonication in isopropanol followed by deionized water and dried under a nitrogen gas stream. For each measurement, a clean coverslip was placed on an oil-immersion objective (Olympus PlanApo N, 60×, 1.42 NA), and a holey silicone gasket was adhered to the top surface. All dilutions and measurements were performed at room temperature (23 ± 2 °C) inside an anaerobic glove box in buffer containing 50 mM Tris (pH 7.2), 150 mM NaCl, and 1.7 mM sodium dithionite. Samples were stored in airtight glass vials with inserts. Immediately prior to measurement, 2 µL of sample was rapidly diluted into 15 µL of buffer using a Hamilton gas-tight syringe to achieve a final concentration of ∼10 nM (unless otherwise noted in the corresponding figure). Videos were recorded for 1 min following sample addition, capturing high-contrast light-scattering events corresponding to individual particle landings on the coverslip. A known mass standard (β-amylase, Sigma A8781-1VL) was used to calibrate contrast-to-mass conversion. Histograms were generated from all detected events within the 1 min acquisition window and fitted using a non-linear least squares approach to a single or multiple Gaussian function, as appropriate for each sample distribution, to determine the mean mass and associated error. All values from the MP measurements are summarized in **Supplemental Table 5**.

