## Supplementary Information for "Cryo-EM structure of a methanogen nitrogenase-PII protein supercomplex"

**Supplementary Figures 1 – 22**

**Supplementary Tables 1 – 5**

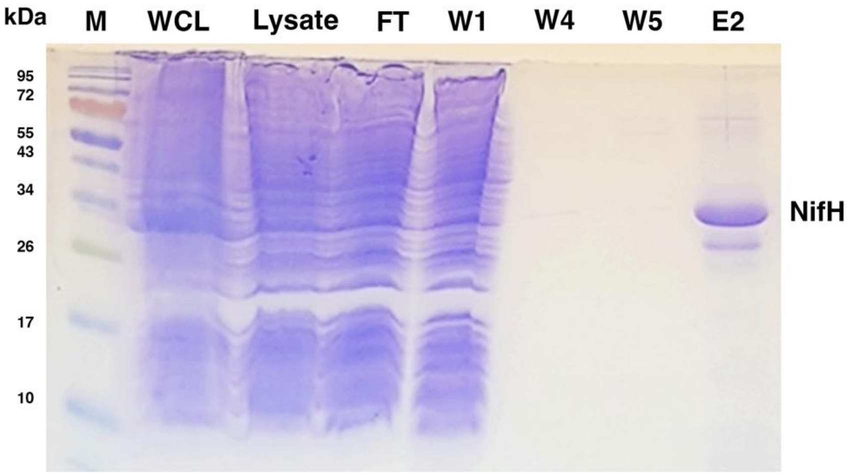


**Supplementary Figure 1. Purification of NifH.** Representative 15% SDS-PAGE gel showing the recombinantly expressed and purified Strep-tagged NifH from *E. coli* SufFeScient cells. M: protein marker; WCL: whole cell lysate; FT: flowthrough; W1–W5: wash fractions; E2: elution fraction 2.

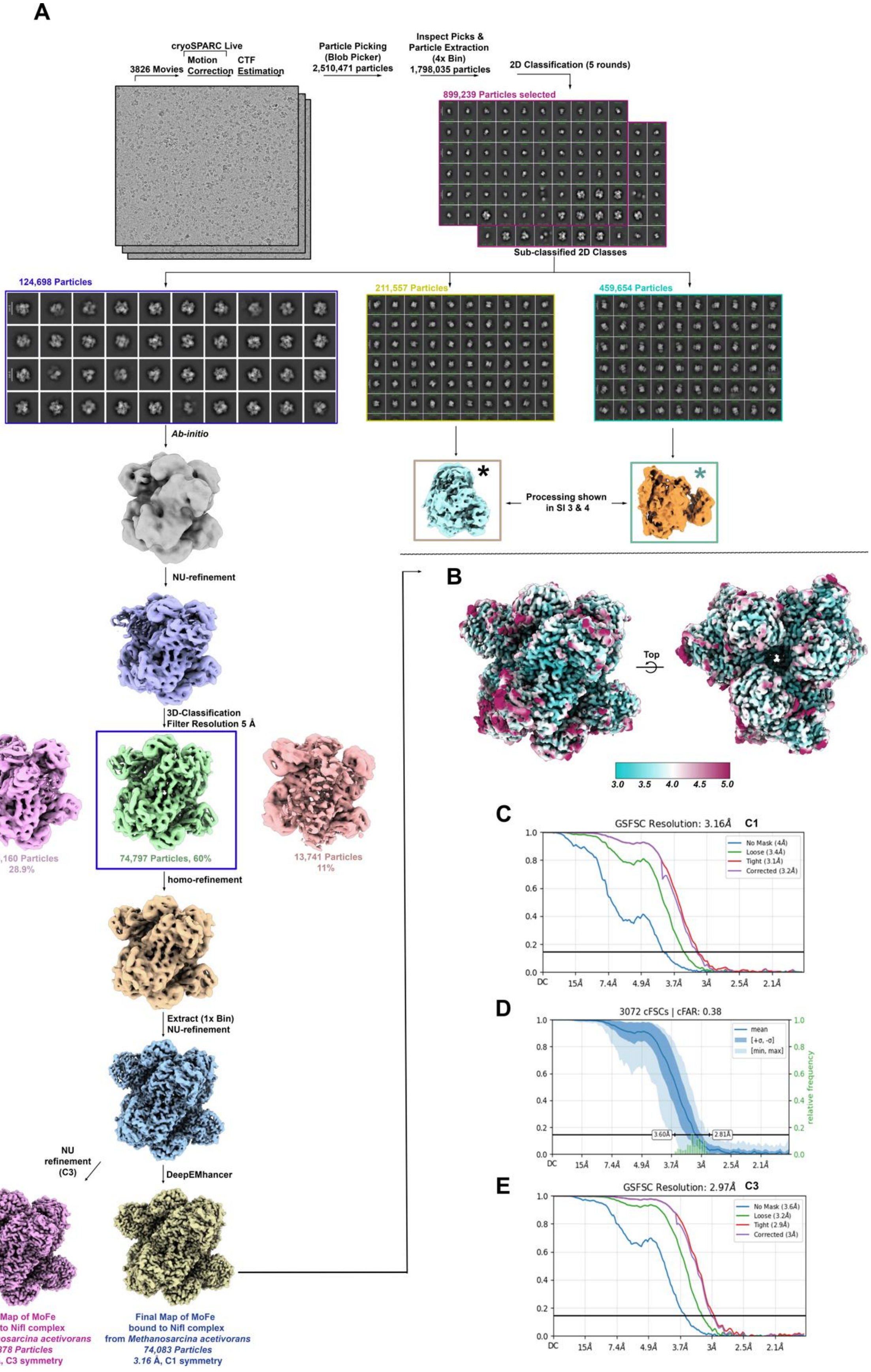

A
cryoSPARC Live
3826 Movies
Motion Correction
CTF Estimation
Particle Picking (Blob Picker) 2,510,471 particles
Inspect Picks & Particle Extraction (4x Bin) 1,798,035 particles
2D Classification (5 rounds)
899,239 Particles selected
Sub-classified 2D Classes
124,698 Particles
211,557 Particles
459,654 Particles
Ab-initio
Processing shown in SI 3 & 4
NU-refinement
3D-Classification Filter Resolution 5 Å
36,160 Particles 28.9%
74,797 Particles, 60%
13,741 Particles 11%
homo-refinement
Extract (1x Bin) NU-refinement
NU refinement (C3)
DeepEMhancer
Final Map of MoFe bound to NifI complex from *Methanosarcina acetivorans* 70,378 Particles 2.97 Å, C3 symmetry
Final Map of MoFe bound to NifI complex from *Methanosarcina acetivorans* 74,083 Particles 3.16 Å, C1 symmetry
B
Top
3.0 3.5 4.0 4.5 5.0
C
GSFSC Resolution: 3.16Å C1
No Mask (4Å)
Loose (3.4Å)
Tight (3.1Å)
Corrected (3.2Å)
DC 15Å 7.4Å 4.9Å 3.7Å 3Å 2.5Å 2.1Å
D
3072 cFSCs | cFAR: 0.38
mean
[+σ, -σ]
[min, max]
relative frequency
3.60Å
2.81Å
DC 15Å 7.4Å 4.9Å 3.7Å 3Å 2.5Å 2.1Å
E
GSFSC Resolution: 2.97Å C3
No Mask (3.6Å)
Loose (3.2Å)
Tight (2.9Å)
Corrected (3Å)
DC 15Å 7.4Å 4.9Å 3.7Å 3Å 2.5Å 2.1Å

**Supplementary Figure 2. Cryo-EM data processing workflow for the (PII-DKKD-PII)$_3$ supercomplex (State-4). (A)** Flowchart of single-particle cryo-EM image processing for the (PII-DKKD-PII)$_3$ supercomplex. Colored boxes indicate particle subsets and corresponding 3D reconstructions selected for further processing. Further processing of the 3D reconstruction marked with a black asterisk (*) is detailed in Supplementary Figure 4, whereas the volume marked with a blue asterisk (*) required additional data collection for further refinement and is detailed in Supplementary Figure 3. **(B)** Local resolution map of the final (PII-DKKD-PII)$_3$ supercomplex reconstruction. **(C)** Gold-standard Fourier shell correlation (FSC) curve for the C1 reconstruction, showing an overall resolution of 3.16 Å, determined using the FSC = 0.143 criterion. **(D)** 3D directional Fourier shell correlation (3D-FSC) plot for the final (PII-DKKD-PII)$_3$ supercomplex reconstruction. The blue curve represents the directional FSC average, and the shaded region indicates the spread among different orientations, demonstrating isotropic resolution distribution. The dashed horizontal line marks the FSC = 0.143 threshold. **(E)** Gold-standard Fourier shell correlation (FSC) curve for the C3-symmetry reconstruction, showing the overall resolution of 2.97 Å, determined using the FSC = 0.143 criterion.

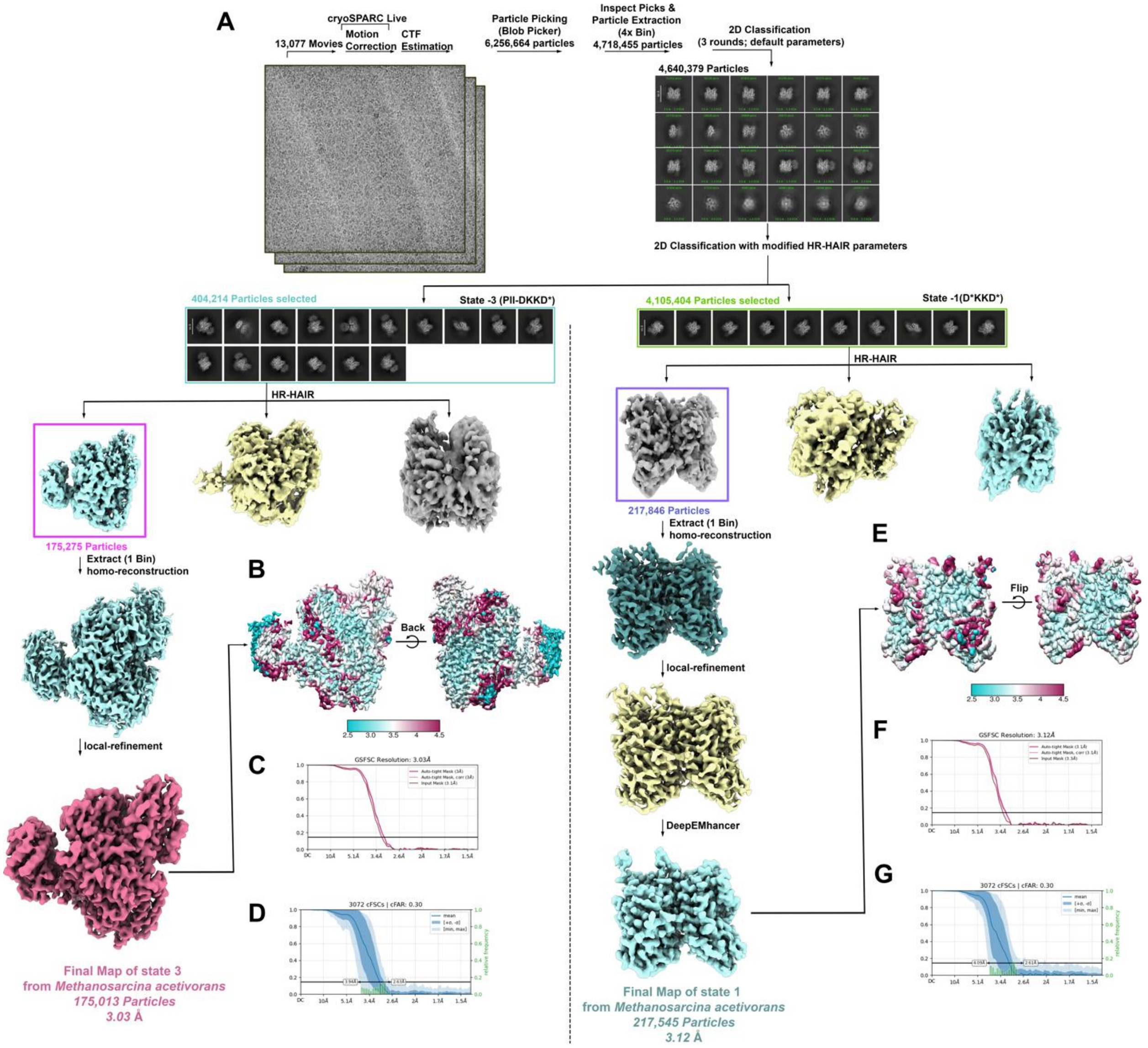


**Supplementary Figure 3. Cryo-EM data processing workflow for the D*KKD* (State-1) and PII-DKKD (State-3) complexes.** (**A**) Flowchart of single-particle cryo-EM image processing for the D*KKD* (right) and PII-DKKD* complexes (left). (**B**) Local resolution map of the final PII-DKKD* complex reconstruction. (**C**) Gold-standard Fourier shell correlation (FSC) curve for the PII-DKKD* complex reconstruction, showing an overall resolution of 3.03 Å, determined using the FSC = 0.143 criterion. (**D**) 3D directional Fourier shell correlation (3D-FSC) plot for the final PII-DKKD* complex reconstruction. (**E**) Local resolution map of the final D*KKD* complex reconstruction. (**F**) Gold-standard Fourier shell correlation (FSC) curve for the D*KKD* complex reconstruction, showing an overall resolution of 3.12 Å, determined using the FSC = 0.143 criterion. (**G**) 3D directional Fourier shell correlation (3D-FSC) plot for the final D*KKD* complex reconstruction.

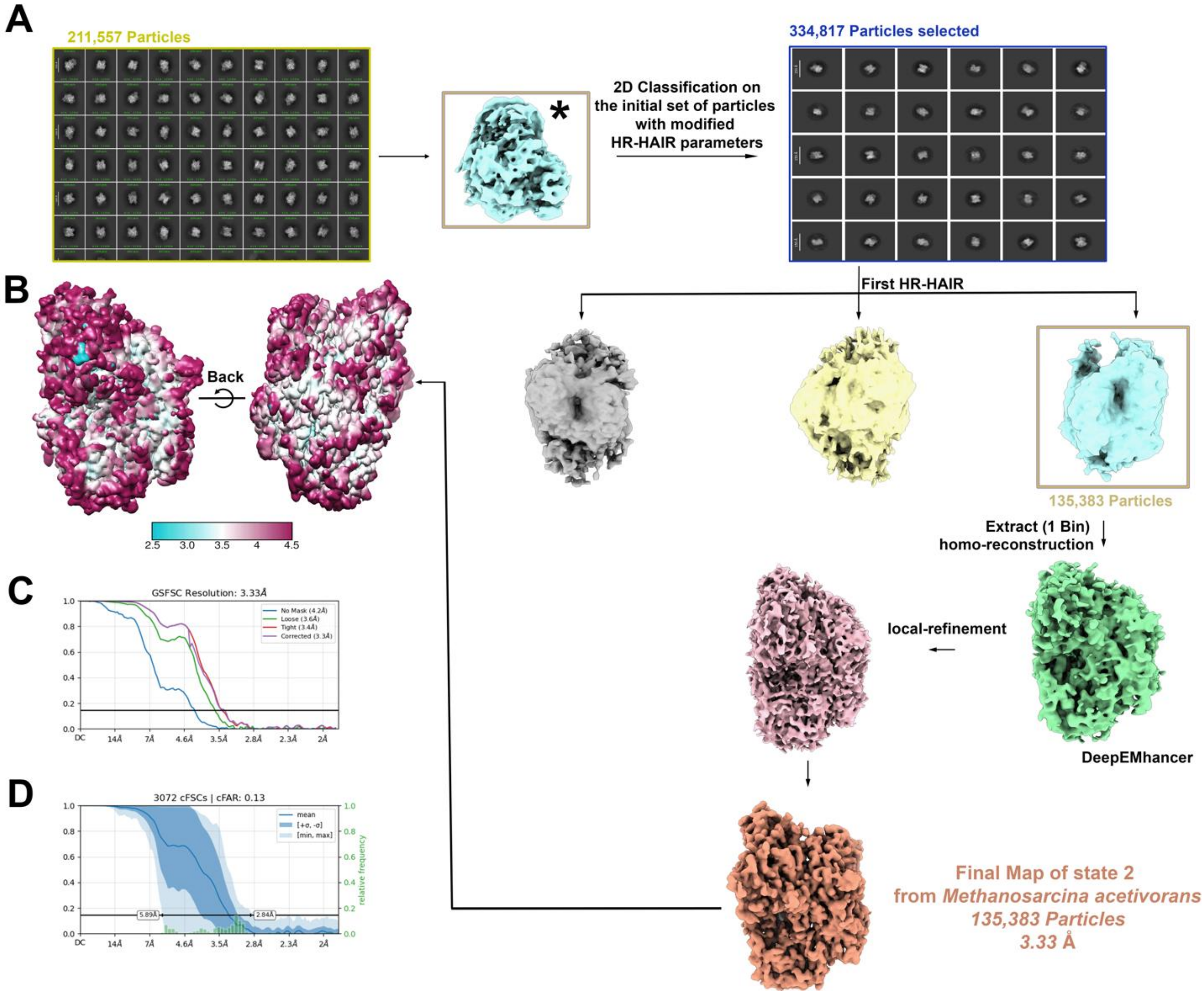


**Supplementary Figure 4. Cryo-EM data processing workflow for the DKK complex (State-2).** (**A**) Flowchart of single-particle cryo-EM image processing for the DKK complex, corresponding to the particle subset marked with a black asterisk (*) in Supplementary Figure 2. (**B**) Local resolution map of the final DKK complex reconstruction. (**C**) Gold-standard Fourier shell correlation (FSC) curve for the DKK complex reconstruction, showing an overall resolution of 3.33 Å, determined using the FSC = 0.143 criterion. (**D**) 3D directional Fourier shell correlation (3D-FSC) plot for the final DKK complex reconstruction.

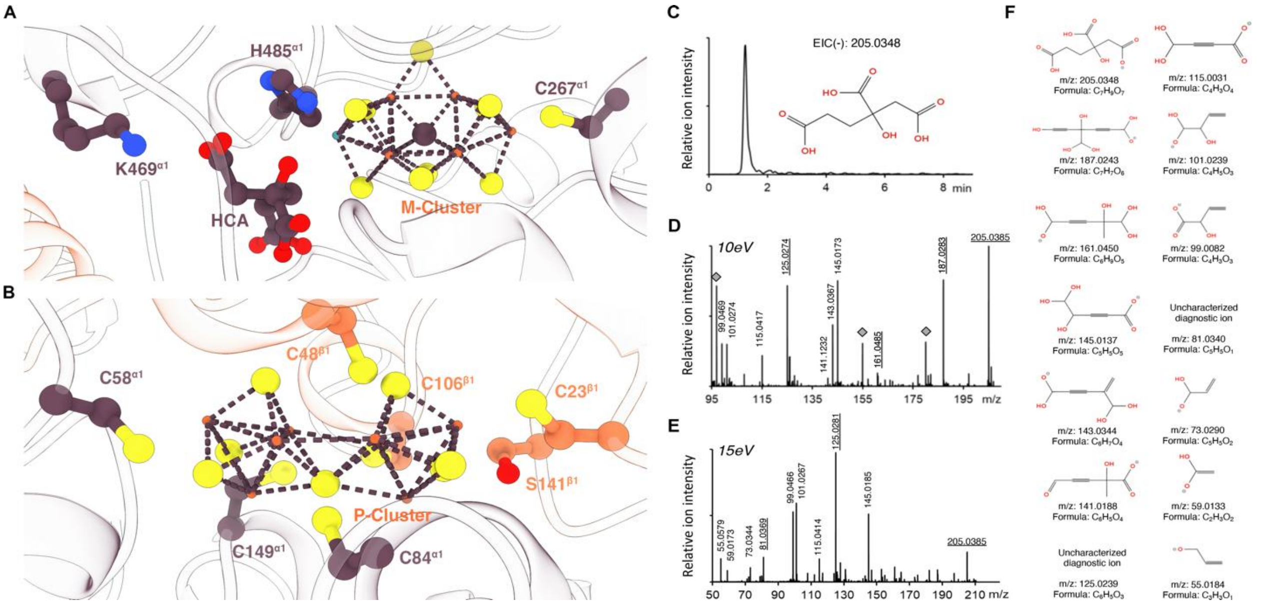


**Supplementary Figure 5. Coordination environment of metal clusters in NifDK from one half of unit 1 and identification of the homocitrate cofactor.** (**A**) Detailed view of the M-cluster coordination, emphasizing the key amino acid residues involved in ligation and stabilization of the cluster. (**B**) Detailed view of the P-cluster coordination, highlighting residues interacting with the iron-sulfur core. (**C**) Represents extracted ion chromatogram (EIC) for m/z=205.0348 which corresponds to the calculated mass of homocitrate ion in a negative mode. (**D-E**) Represents the fragmentation pattern of m/z=205.0348 collected at two different collision energies: 10 eV and 15eV, respectively. Annotated peaks represent identified fragments; underlined values represent diagnostic ions, specific only to the homocitrate molecule. m/z peaks denoted with gray diamonds in (**D**) represent signals from instrument background contaminants. (**F**) The ionic structures and their m/z values are shown.

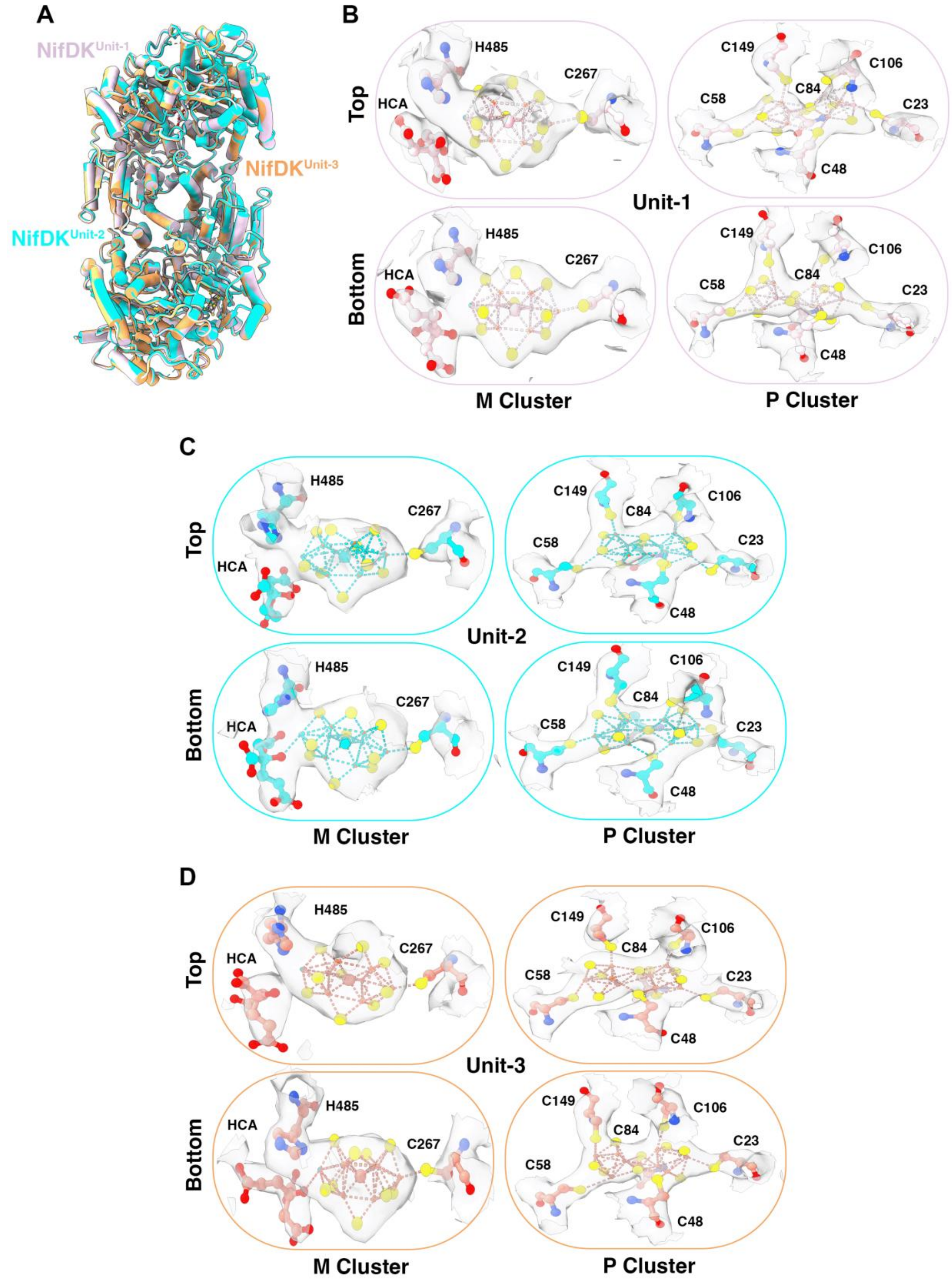


**Supplementary Figure 6. Structural features of the three NifDK heterotetramers in the supercomplex.** (**A**) Overlay of the three NifDK heterotetramers from the three units as shown in cylinder/stubs representation, demonstrating their overall structural similarity. (**B-D**) Electron density maps corresponding to the M-cluster and P-cluster for unit 1, 2 and 3 respectively, with coordinating residues displayed for improved visualization.

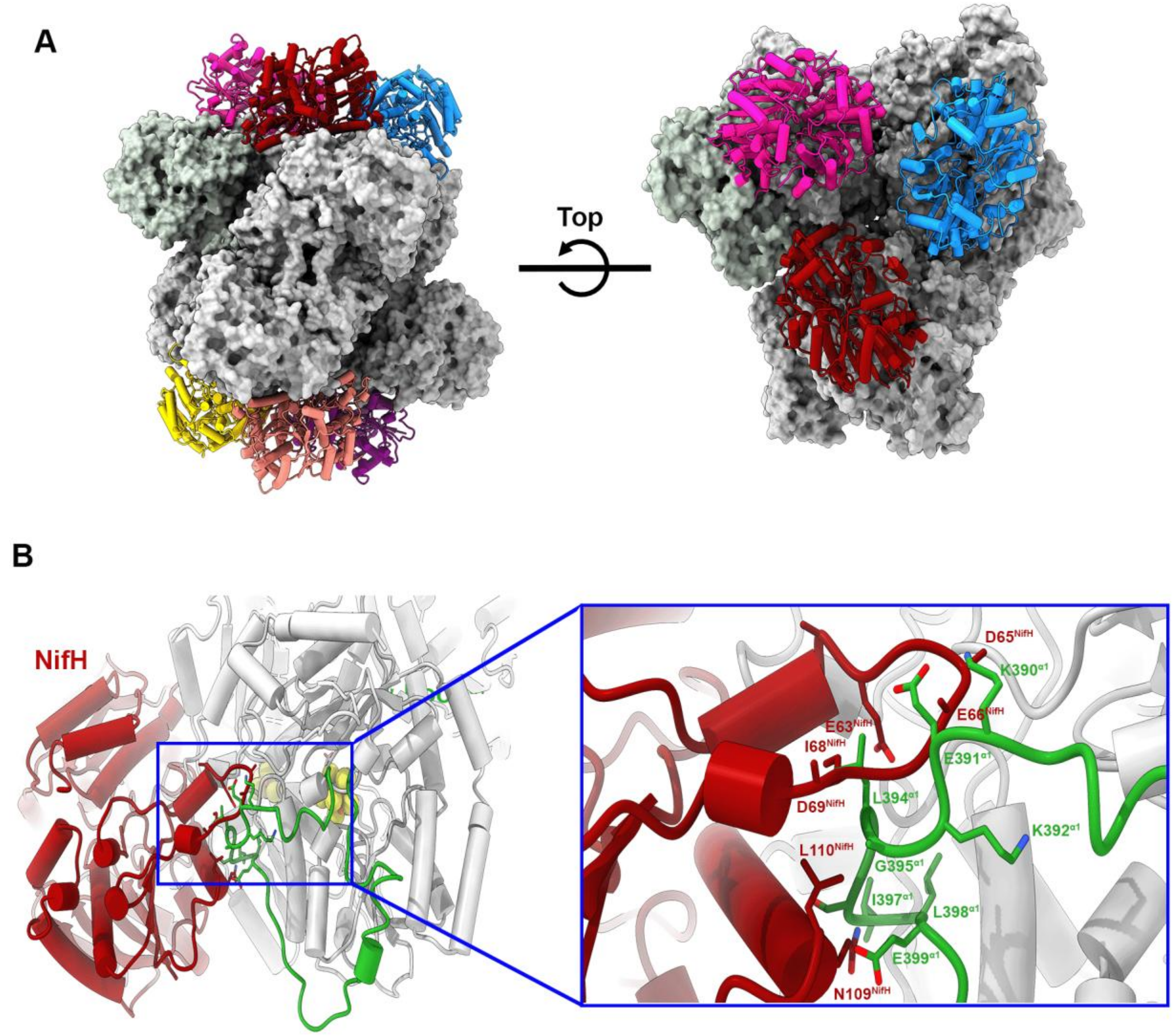


**Supplementary Figure 7. Manual docking of NifH onto the NifDK supercomplex and identification of potential interaction residues.** (**A**) NifH was manually docked onto the NifDK in the supercomplex after removal of the NifI inhibitor complex. The position of NifH (PDB:6NZJ) on NifDK was guided by the structure of *Av* nitrogenase (PDB:4WZB). All the NifDK proteins in the supercomplex can engage a respective NifH without steric hinderance. (**B**) Close up view of the interface highlighting probable residues in the α-loop of NifD involved in interactions with NifH, including a network of salt bridges, hydrogen bonds, and hydrophobic contacts that may stabilize complex assembly.

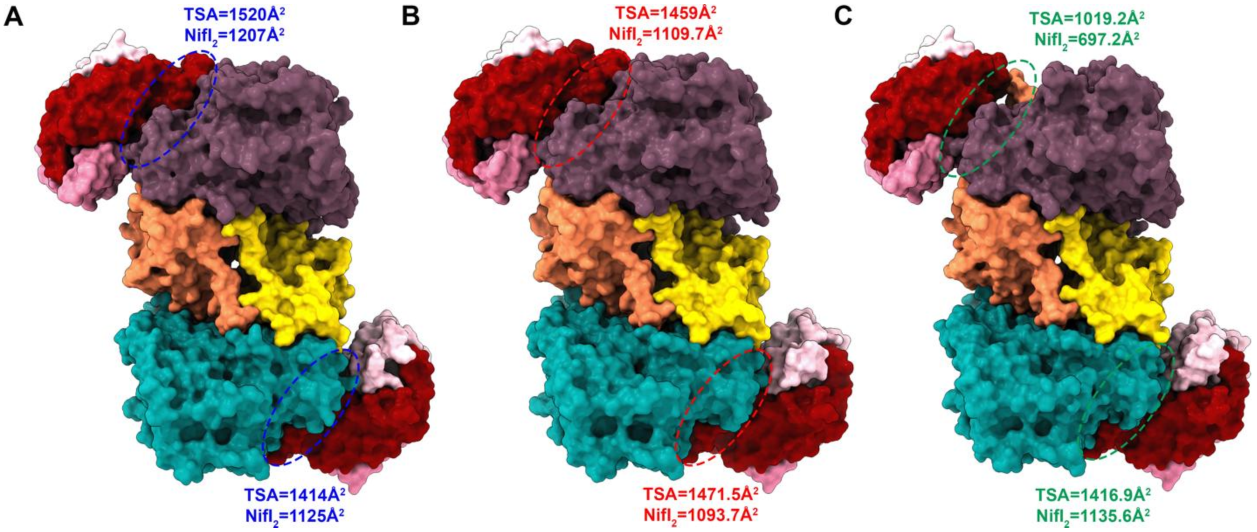


**Supplementary Figure 8. Differential buried surface area between $NifI_2$ within the NifI complex and NifDK across the supercomplex units** (**A**) Unit 1, (**B**) Unit 2, and (**C**) Unit 3. Variations in the buried surface area at the interface are shown with the interface between the NifI complex and NifDK highlighted by dotted ovals in distinct colors for each unit. TSA denotes total surface area.

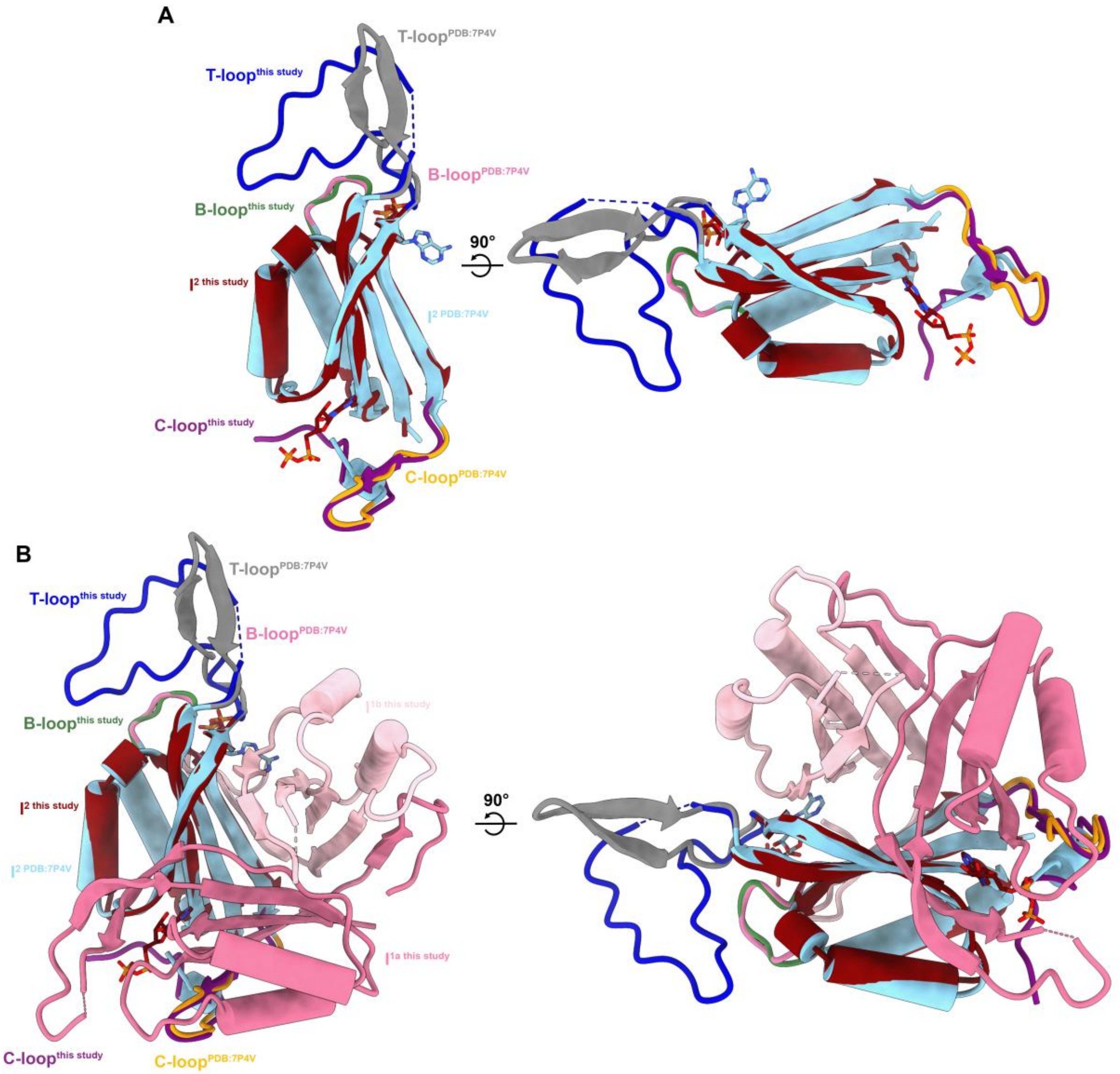


**Supplementary Figure 9. Structural comparison of Glnk and NifI$_2$ from the NifI complex**. (**A**) Overlay of NifI$_2$ with Glnk1 from *Methanothermococcus thermolithotrophicus* (PDB: 7P4V), with the T-, B- and C-loops labeled. Differences in the T- and C-loops between Glnk1 and NifI2 are highlighted. (**B**) Overlay of the entire NifI complex with one Glnk1 monomer, emphasizing the presence of the T-, B-, and C-loops in the NifI complex. 2-oxoglutarate (2OG) and ADP are shown to denote the binding site for ligands.

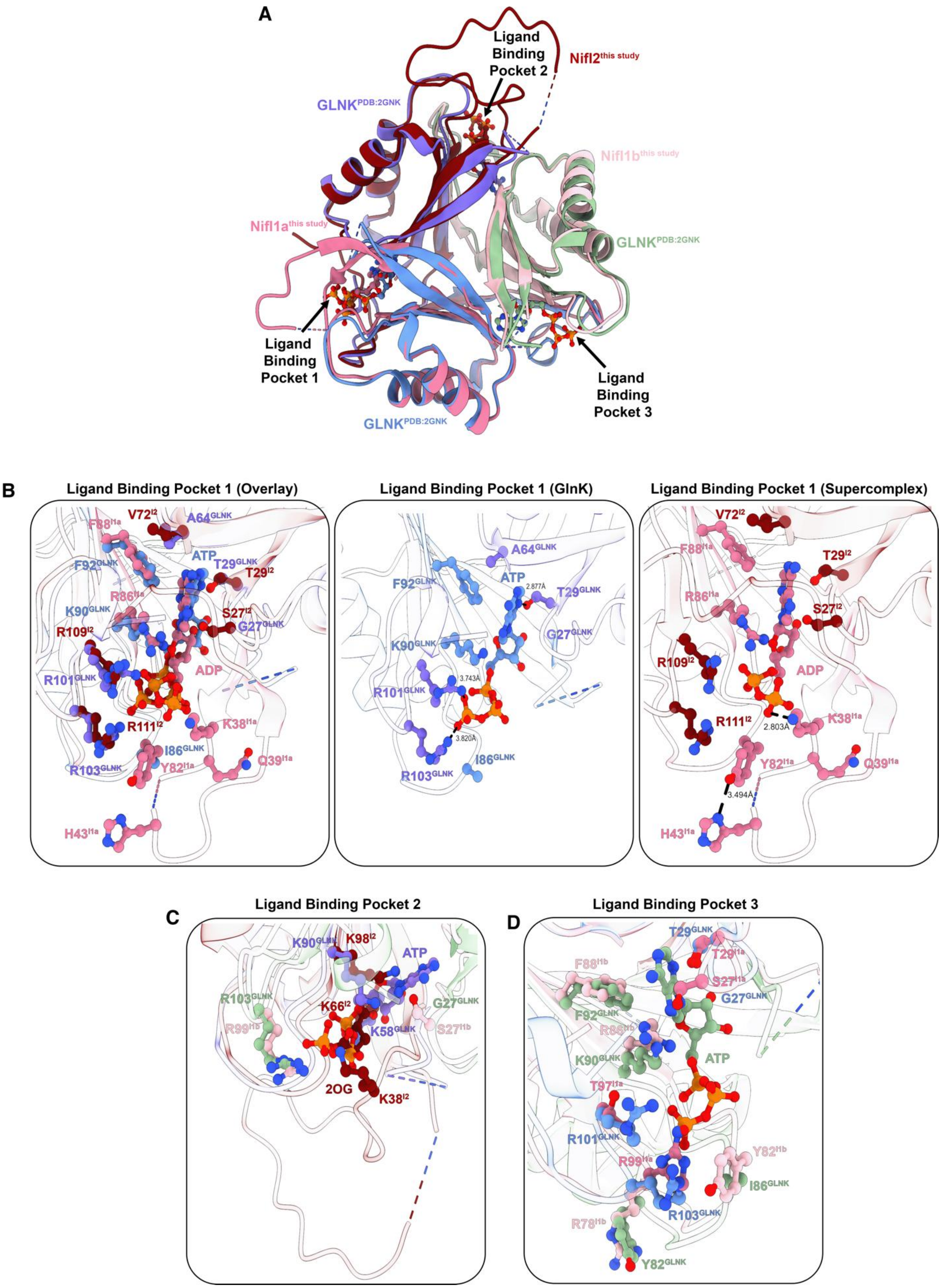
A
Ligand Binding Pocket 2
NifI2this study
GLNKPDB:2GNK
NifI1bthis study
NifI1athis study
GLNKPDB:2GNK
Ligand Binding Pocket 1
Ligand Binding Pocket 3
GLNKPDB:2GNK
B
Ligand Binding Pocket 1 (Overlay)
V72I2
A64GLNK
F88I1a
ATP
F92GLNK
T29GLNK
T29I2
R86I1a
K90GLNK
S27I2
R109I2
G27GLNK
ADP
R101GLNK
R111I2
K38I1a
I86GLNK
R103GLNK
Y82I1a
Q39I1a
H43I1a
Ligand Binding Pocket 1 (GlnK)
A64GLNK
ATP
2.877Å
F92GLNK
T29GLNK
G27GLNK
K90GLNK
3.743Å
R101GLNK
3.820Å
R103GLNK
I86GLNK
Ligand Binding Pocket 1 (Supercomplex)
V72I2
F88I1a
T29I2
R86I1a
S27I2
R109I2
ADP
R111I2
2.803Å
K38I1a
Y82I1a
Q39I1a
3.494Å
H43I1a
C
Ligand Binding Pocket 2
K90GLNK
K98I2
ATP
R103GLNK
K66I2
G27GLNK
R99I1b
K58GLNK
S27I1b
2OG
K38I2
D
Ligand Binding Pocket 3
T29GLNK
F88I1b
T29I1a
S27I1a
G27GLNK
F92GLNK
R86I1b
ATP
K90GLNK
T97I1a
R101GLNK
Y82I1b
R99I1a
I86GLNK
R103GLNK
R78I1b
Y82GLNK

**Supplementary Figure 10. Comparison of ligand binding pockets and interactions between Glnk and the NifI complex from the supercomplex.** (**A**) Overlay of the Glnk structure (PDB: 2GNK) with the NifI complex, highlighting the three ligand binding pockets. (**B**) ADP-bound pocket 1 in the NifI complex (Unit 1) is overlaid with the ATP-bound pocket in the Glnk structure. Key residues that interact with their respective ligand are highlighted for both. For clarity, ADP and ATP are shown in separate panels within their respective binding pockets. (**C**) Ligand binding pocket 2 in the NifI complex bound to 2-oxoglutarate (2OG) is compared against the ATP-bound Glnk. Ligand coordinating residues are. (**D**) Ligand binding pocket 3 in the NifI complex supercomplex structure lacking a bound ligand is compared against the ATP-bound pocket in Glnk.

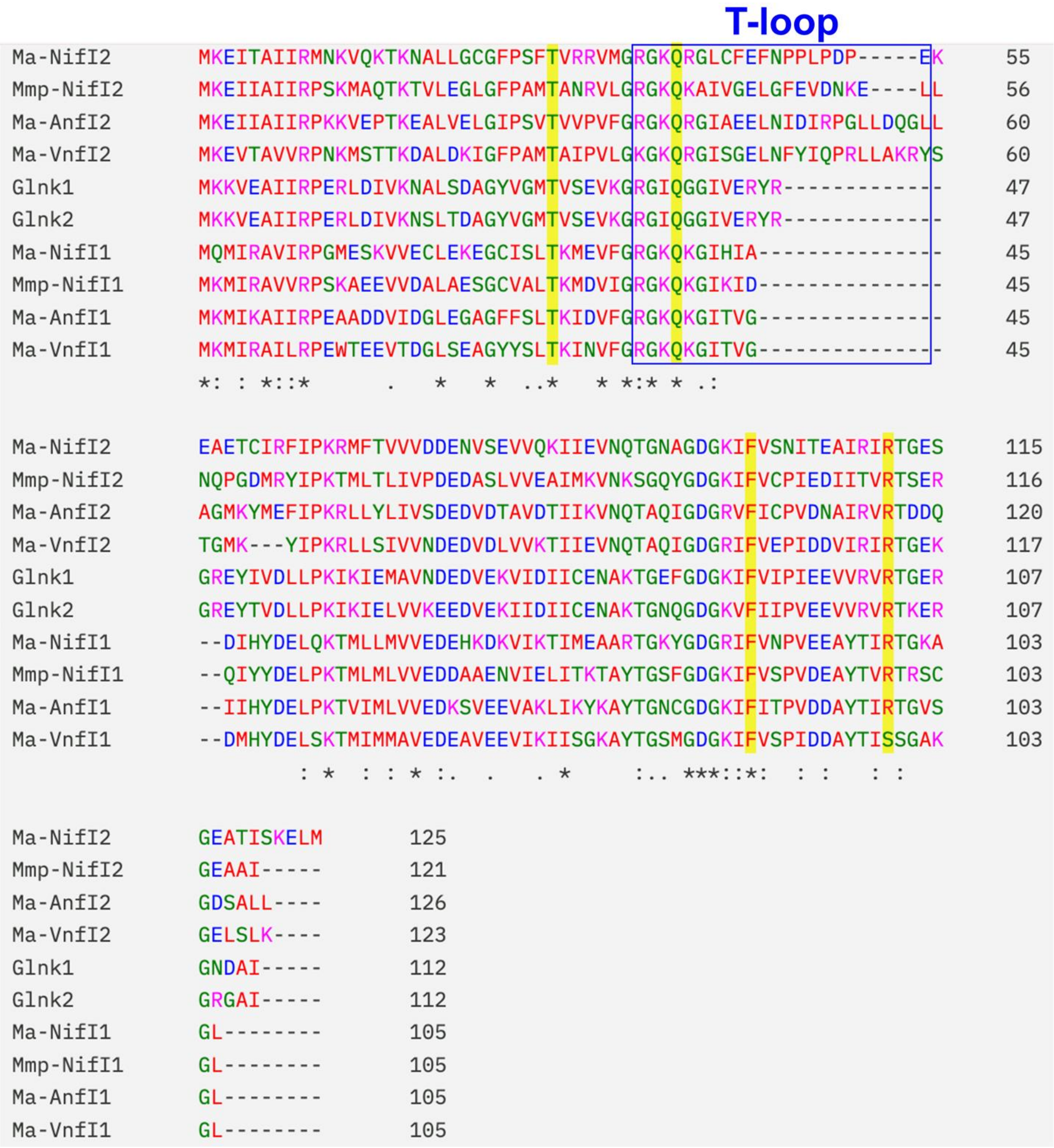


**Supplementary Figure 11. Conservation of ligand binding residues in NifI.** Sequence alignment of PII proteins including Glnk1, Glnk2, NifI$_1$ and NifI$_2$ protein from *Methanosarcina acetivorans* (*Ma*) and *Methanococcus maripaludis* (*Mmp*) highlighting the T-loop and the conserved residues across the species in the ligand binding pocket.

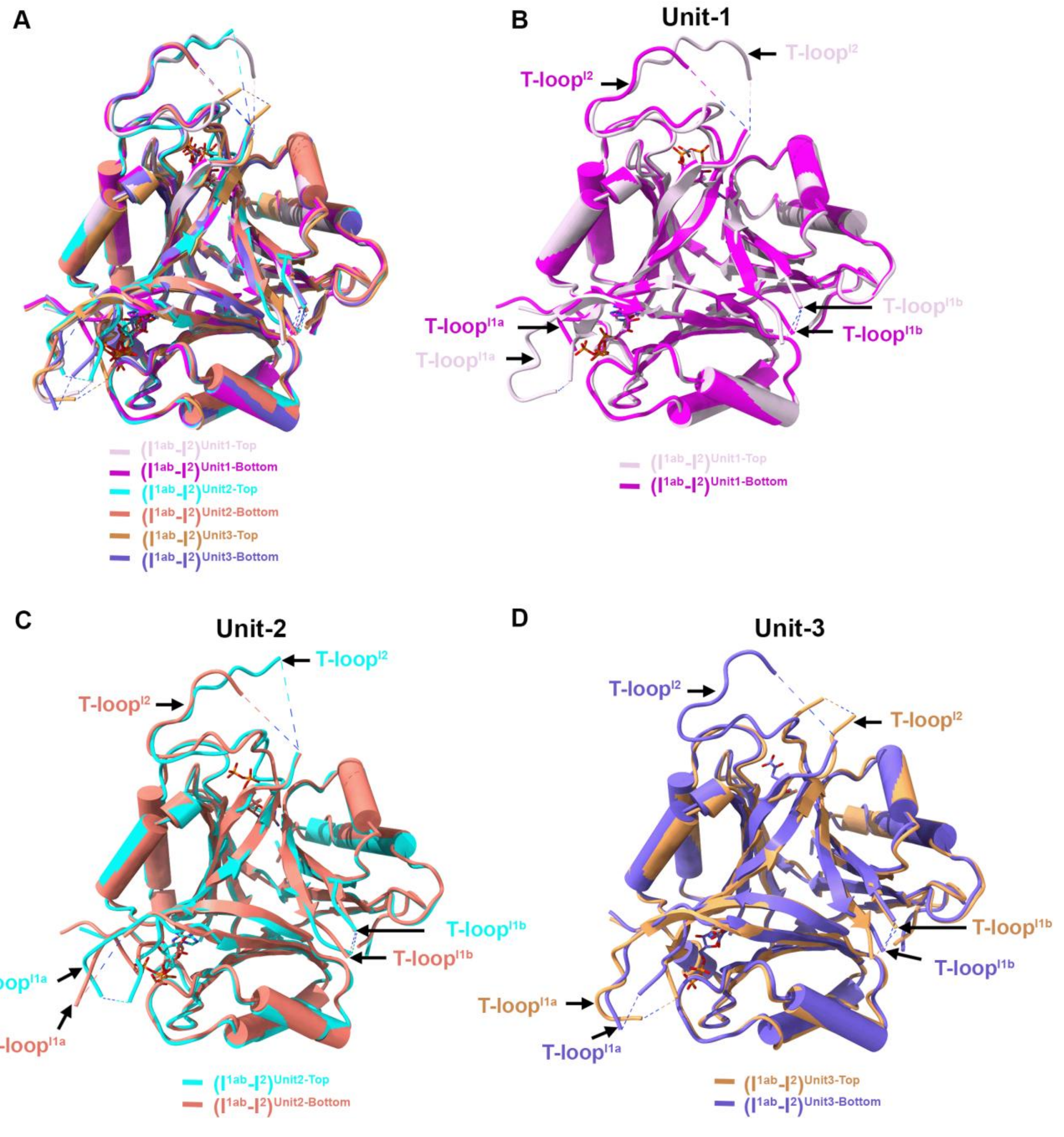


**Supplementary Figure 12. Overlay of the NifI complexes within the supercomplex** (**A**) Overlay of all NifI complexes present in the supercomplex structure. (**B**) Overlay of the top and bottom NifI complexes from unit 1. (**C**) Overlay of the top and bottom NifI complexes from unit 2. (**D**) Overlay of the top and bottom NifI complexes from unit 3. The T-loops are marked.

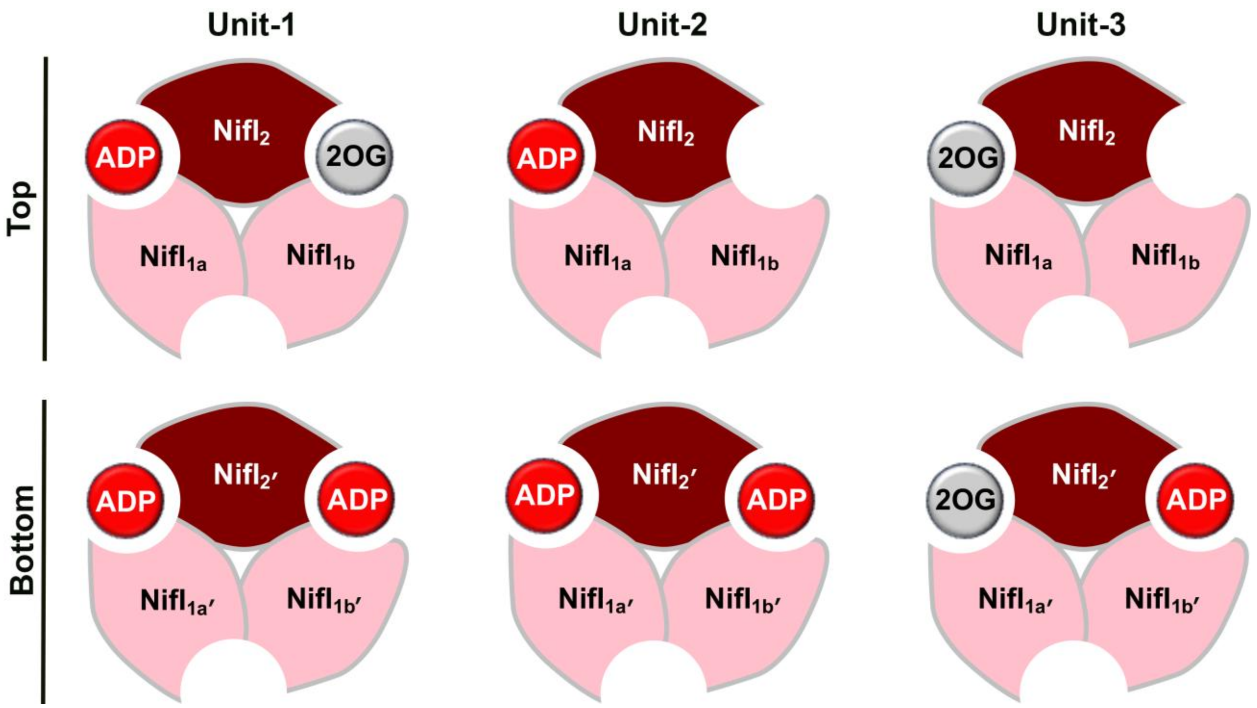


**Supplementary Figure 13. Schematic of the ligand occupancies in the NifI proteins bound in the supercomplex**. Two of the NifI complexes in the supercomplexs (Unit 1′ and Unit 2′) are bound to two ADP molecules. Similarly, two of the NifI complexes are bound to one ADP and 2OG each in the supercomplex (Unit 1 and Unit 3′). ADP or 2OG binding is captured only between the $NifI_1$-$Nif_2$ and $NifI_2$-$Nif_1$ interfaces and never in the $NifI_1$-$NifI_2$ interface in the supercomplex.

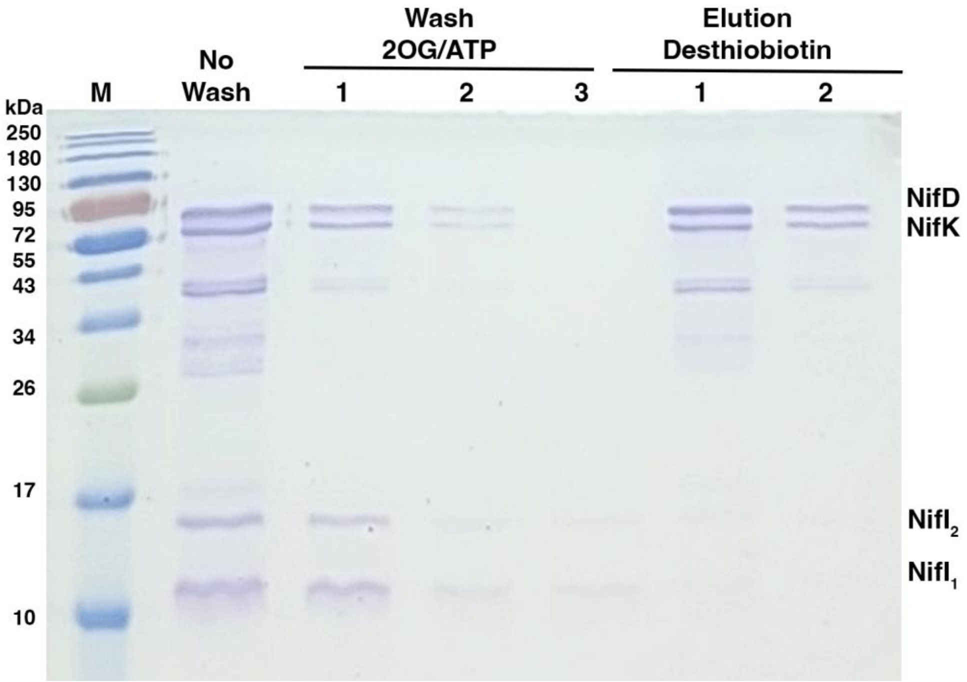


**Supplementary Figure 14. The effect of 2OG and ATP on complex formation.** Representative SDS-PAGE gel illustrating the purification of NifDK complex after washing resin-bound strep-NifD with 2OG and $Mg^{2+}$- ATP (10 mM each). 2OG and ATP effectively remove $NifI_1$ and $NifI_2$ from bound strep-NifD, as evidenced by their absence in the elution fraction and presence in the wash fractions.

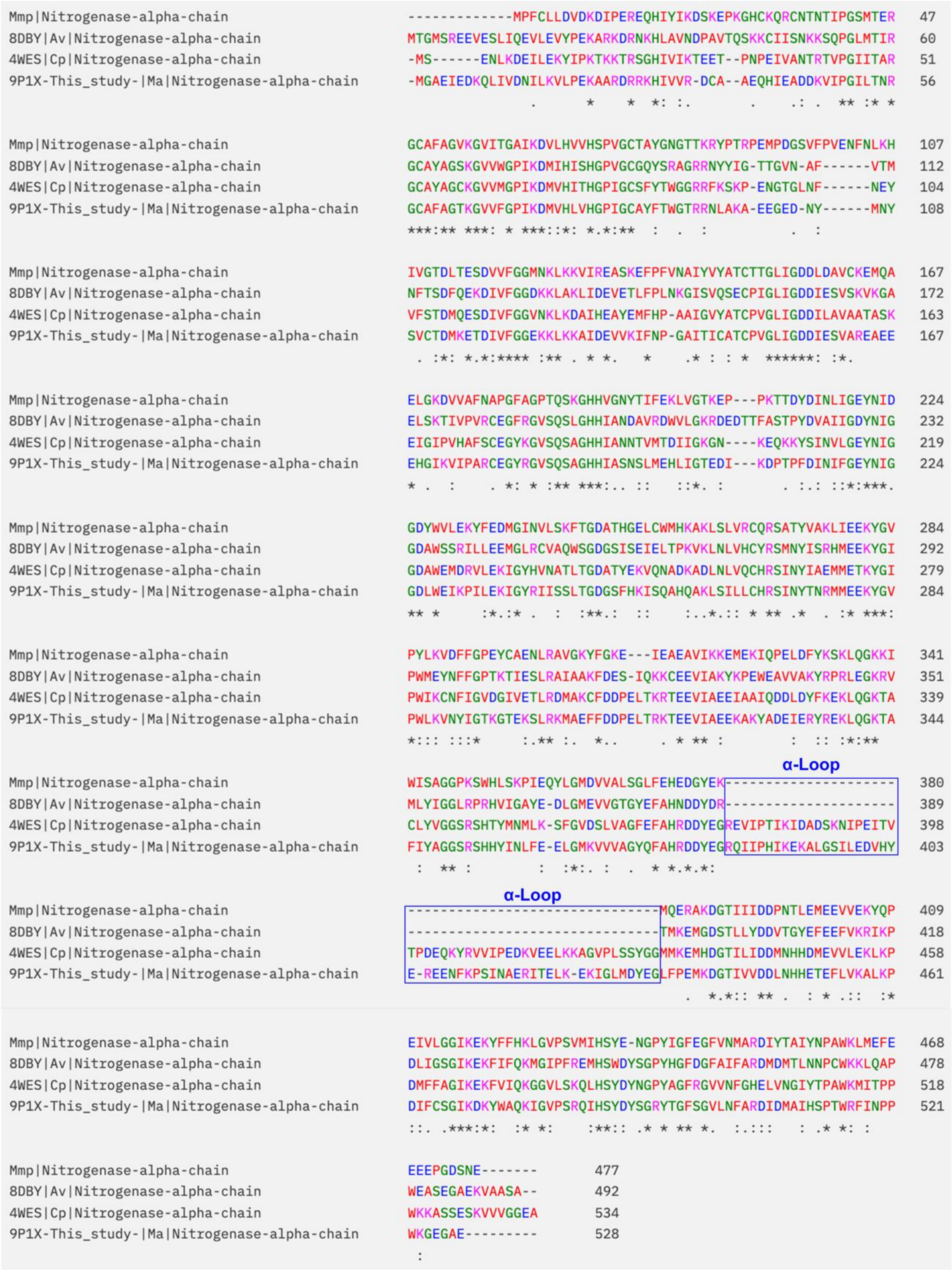


**Supplementary Figure 15. Conservation of α-loop in *Methanosarcina acetivorans*.** Sequence alignment of NifD proteins from *Methanococcus maripaludis* (*Mmp*), *Azotobacter vinelandii* (*Av*), *Clostridium pasteurianum* (*Cp*) and *Methanosarcina acetivorans* (*Ma*), *highlighting* the α-loop present in *Ma* and *Cp.*

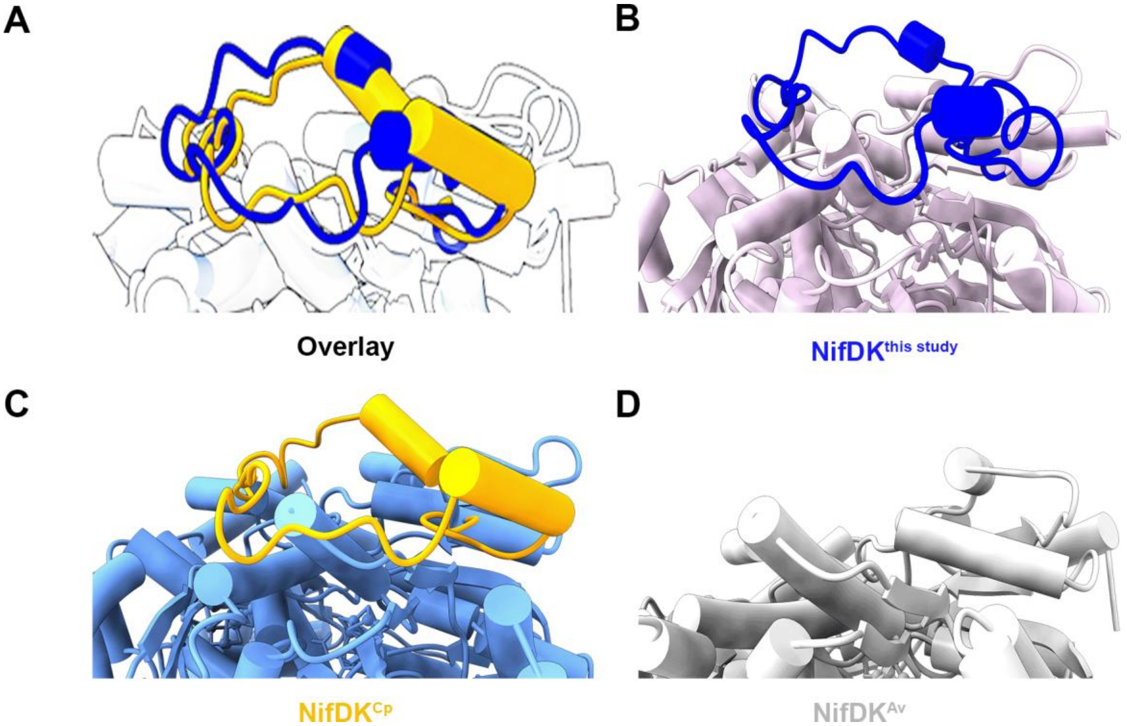


**Supplementary Figure 16. Zoomed-in view of key loops highlighted in Figure 7G for clearer visualization. (A)** Overlay of the α- and α′-loops from supercomplex in this study (blue) with those from *C. pasteurianum* (gold, PDB: 8DBY) and *A. vinelandii* (grey, PDB: 4WES), illustrating conformational diversity of the loops. **(B–D)** Non-superimposed views of the three loops are shown for clarity.

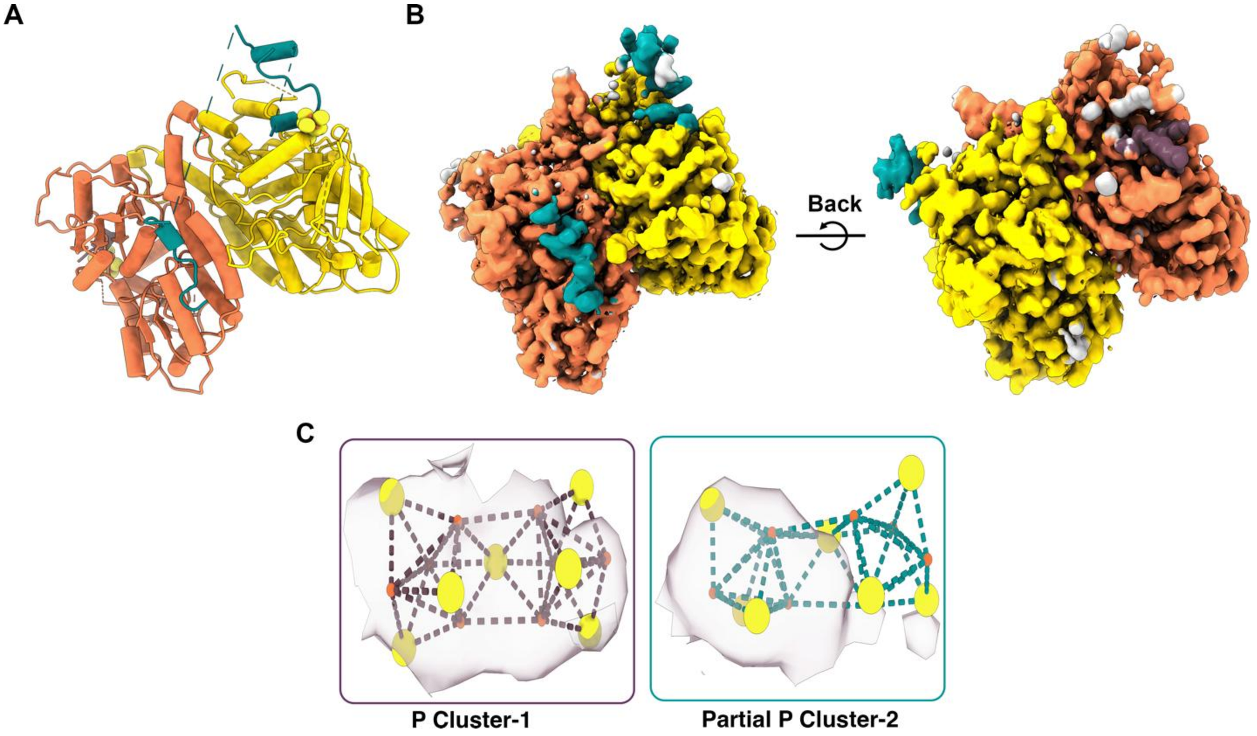


**Supplementary Figure 17. Structural features of the D*KKD* complex (State-1). (A)** Overall structure of the D*KKD* complex. **(B)** Corresponding electron density map, with additional density from dynamic NifD chains (grey) not included in the modeled structure. **(C)** Carved electron density for the P-clusters observed on either side of the complex.

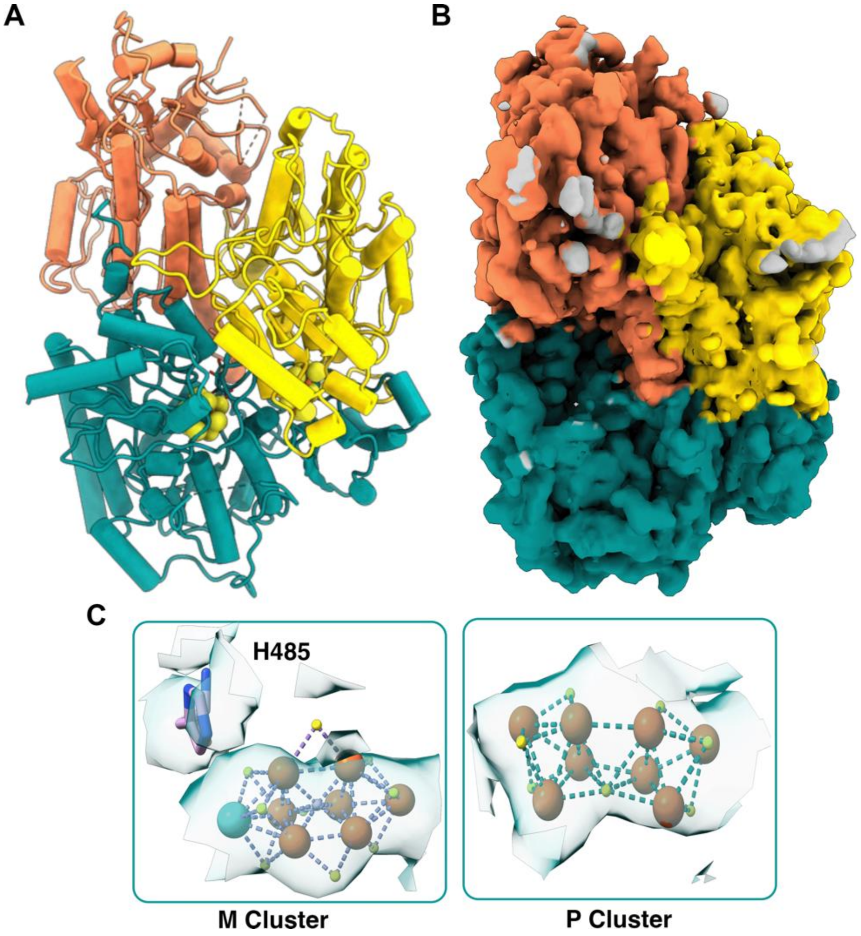


**Supplementary Figure 18. Structural features of the DKK complex (State-2). (A)** Overall structure of the DKK complex. **(B)** Corresponding electron density map, showing additional density from dynamic NifD chains (grey) not included in the modeled structure. Although some density for the P-cluster on the opposite side is visible, it was not modeled to ensure confidence in model building. **(C)** Carved electron density for the M- and the P-cluster observed in the complex.

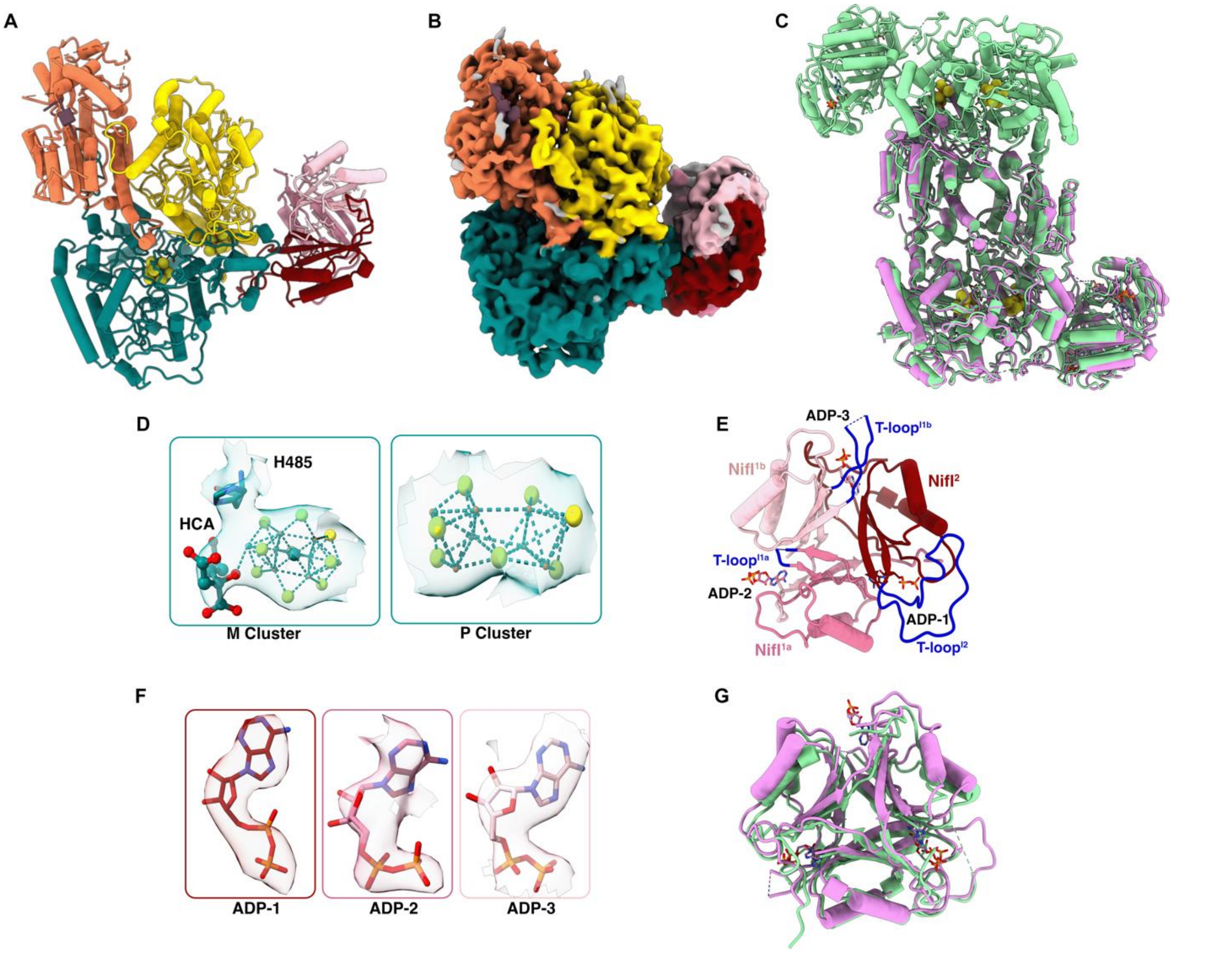


**Supplementary Figure 19. Structural features of the PII-DKKD* complex (State-3). (A)** Overall structure of the PII-DKKD* complex. **(B)** Corresponding electron density map, showing additional density from dynamic NifD chains (grey) not included in the modeled structure. **(C)** Overlay of the one of the units from the supercomplex (green) with this structure (magenta), highlighting overall structural similarity and the missing NifD unit. **(D)** Carved electron density for the M- and P-clusters observed in the PII-DKKD* complex. **(E)** Close-up view of the PII complex extracted from this structure, with T-loops highlighted in blue. Individual NifI subunits are shown in distinct colors, consistent with the color scheme used throughout the manuscript. Bound ADP molecules associated with each subunit are depicted as sticks. **(F)** Carved density for ADP observed in the ligand-binding pockets for all three sites of the PII-DKKD* complex. **(G)** Overlay of one of the PII complex from the supercomplex (green) with this structure (magenta), showing overall similarity and differences in the loop region.

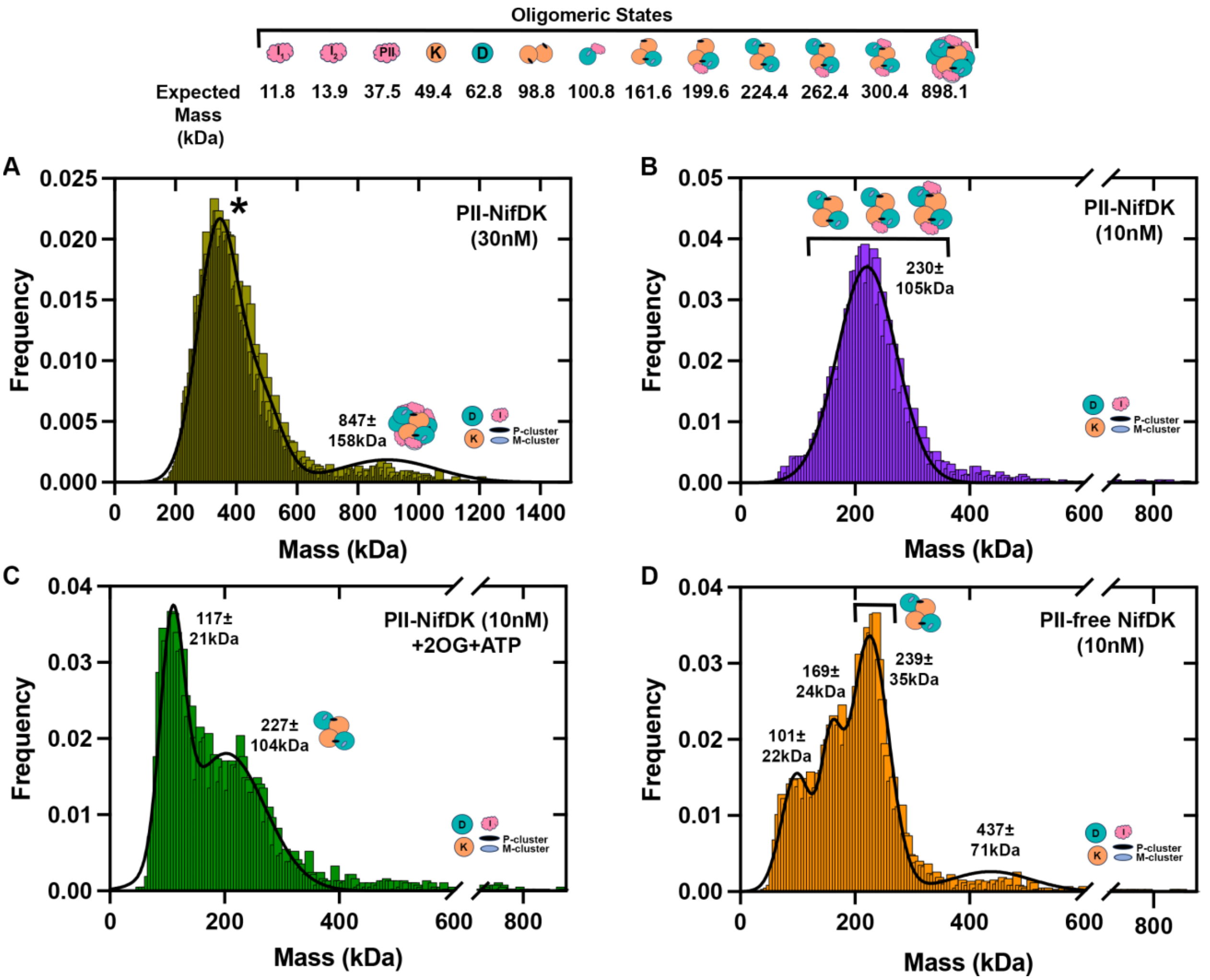


**Supplementary Figure 20. Mass photometry (MP) analysis of NifDK complexes.** MP of the native Strep-NifD pull down sample was measured at **(A)** 30 nM or **(B)** 10 nM. Higher concentrations were required to capture the supercomplex species (calculated mass 898 kDa). But, due to the high particle count for the major species (*), the mass interpretation is overestimated. The lower concentration measurement in panel B generates appropriate particle numbers for the major species and reflects the NifDK tetramer (calculated mass 225 kDa). However, the distribution is broad and likely reflects both PII-bound and PII-free NifDK tetramers. **(C)** The native Strep-NifD pull down sample was mixed with 2OG (0.1 mM) and ATP-$Mg^{2+}$ (0.1 mM) and MP analysis shows presence of the NifDK tetramer along with transitions to a wide range of smaller sized complexes. **(D)** The PII-free NifDK sample was analyzed by MP. The NifDK tetramer is the major species along with both larger and smaller sized complexes.

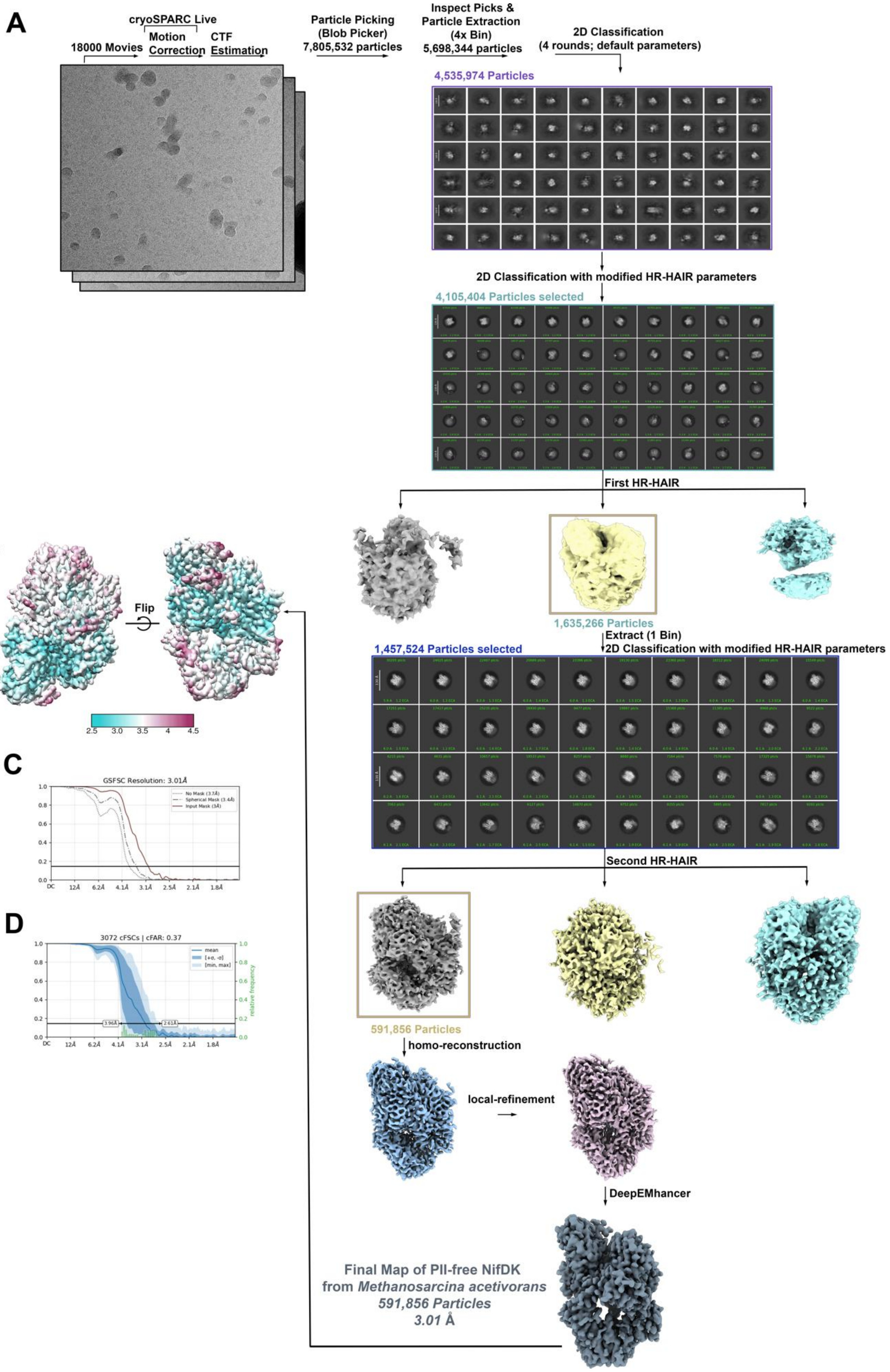
A
cryoSPARC Live
18000 Movies
Motion Correction
CTF Estimation
Particle Picking (Blob Picker)
7,805,532 particles
Inspect Picks & Particle Extraction (4x Bin)
5,698,344 particles
2D Classification (4 rounds; default parameters)
4,535,974 Particles
2D Classification with modified HR-HAIR parameters
4,105,404 Particles selected
First HR-HAIR
1,635,266 Particles
Extract (1 Bin)
2D Classification with modified HR-HAIR parameters
1,457,524 Particles selected
Second HR-HAIR
591,856 Particles
homo-reconstruction
local-refinement
DeepEMhancer
Final Map of PII-free NifDK from Methanosarcina acetivorans
591,856 Particles
3.01 Å
B
Flip
2.5
3.0
3.5
4
4.5
C
GSFSC Resolution: 3.01Å
No Mask (3.7Å)
Spherical Mask (3.4Å)
Input Mask (3Å)
D
3072 cFSCs | cFAR: 0.37
mean
relative frequency
3.96Å
2.61Å

**Supplementary Figure 21. Cryo-EM data processing workflow for the PII-free NifDK complex.** **(A)** Flowchart of single-particle cryo-EM image processing for the PII-free NifDK complex. **(B)** Local resolution map of the final PII-free NifDK complex reconstruction. **(C)** Gold-standard Fourier shell correlation (FSC) curve for the PII-free NifDK complex reconstruction, showing an overall resolution of 3.01 Å, determined using the FSC = 0.143 criterion. **(D)** 3D directional Fourier shell correlation (3D-FSC) plot for the final PII-free NifDK complex reconstruction.

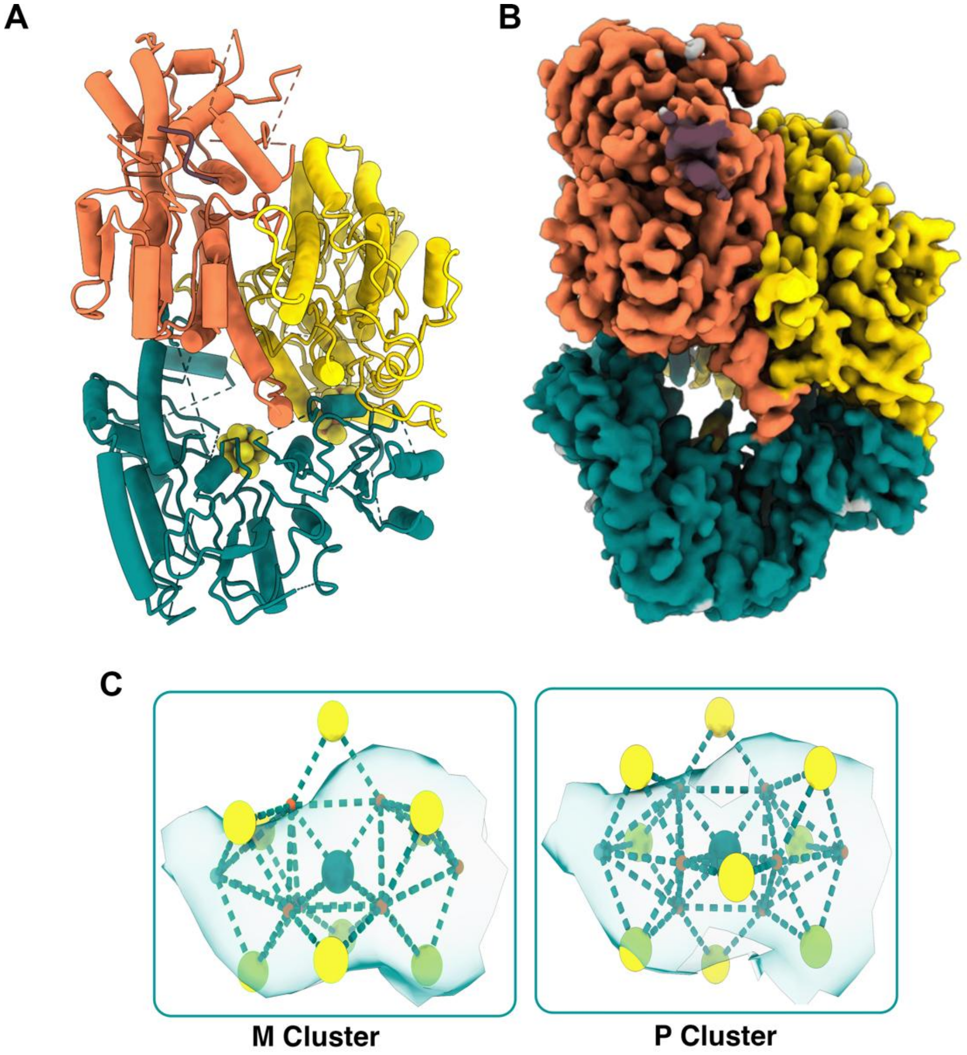


**Supplementary Figure 22. Structural features of the PII-free D#KKD* complex (State-5). (A)** Overall structure of the PII-free D#KKD* complex. **(B)** Corresponding electron density map, showing additional density from dynamic NifD chains (eggplant) modelled in the structure. **(C)** Carved electron density for the M- and the P-cluster observed in the complex.

**Supplementary Table 1.** Cryo-EM data collection and model statistics of the *M. acetivorans* NifDK bound to NifI supercomplex in C1 and C3 symmetry.

| | **NifDK bound to NifI Supercomplex (C1)** (EMDB- 71144); (PDB-9P1X) | **NifDK bound to NifI Supercomplex (C3)** (EMDB- 76263); (PDB-12AH) |
|---|---|---|
| **Data Collection** | TFS Glacios FEI FALCON IV | TFS Glacios FEI FALCON IV |
| Magnification | 150,000 | 150,000 |
| Voltage (kV) | 200 | 200 |
| Spherical Aberration (mm) | 2.7 | 2.7 |
| Electron Exposure ($e^-$/ Å$^2$) | 52 | 52 |
| Defocus range (mm) | -0.1 to -2.4 | -0.1 to -2.4 |
| Pixel size (Å, Physical/Digital) | 0.928 | 0.928 |
| Movies | 3826 | 3826 |
| **Map Statistics and Post-Processing** | | |
| Symmetry imposed | C1 | C3 |
| Map Resolution (Å) | 3.16 | 2.97 |
| FSC threshold | 0.143 | 0.143 |
| **Model Composition** | | |
| Chains | 30 | 43 |
| Non-hydrogen atoms | 60056 | 60643 |
| Protein residues | 7690 | 7724 |
| Ligands | ADP:7<br>AKG:3<br>CLF:6<br>HCA:6<br>ICS:6 | ADP:9<br>AKG:2<br>CLF:6<br>HCA:6<br>ICS:6 |
| Water | 0 | 0 |
| Bonds (RMSD)<br>Length (Å) (#>4s)<br>Angles (º) (#>4s) | <br>0.003 (0)<br>0.542 (21) | <br>0.005 (0)<br>0.644 (0) |
| **Validation** | | |
| Molprobity Score | 2.40 | 2.18 |
| Ramachandran plot<br>Favored (%)<br>Allowed (%)<br>Outliers (%) | <br>94.74<br>5.26<br>0.00 | <br>95.04<br>4.95<br>0.01 |

**Supplementary Table 2.** Cryo-EM data collection and model statistics of the different smaller states of *M. acetivorans* NifDK.

| | **State-1**<br>**D*KKD***<br>(EMDB- 76071);<br>(PDB-11UY) | **State-2**<br>**DKKD***<br>(EMDB- 76025);<br>(PDB-11SX) | **State-3**<br>**PII-DKKD***<br>(EMDB- 75852);<br>(PDB-11MY) | **State-5**<br>**D#KKD***<br>(EMDB- 76217);<br>(PDB-11ZK) |
|---|---|---|---|---|
| **Data Collection** | TFS Krios FEI FALCON IV | TFS Glacios FEI FALCON IV | TFS Krios FEI FALCON IV | TFS Krios FEI FALCON IV |
| Magnification | 130,000 | 150,000 | 130,000 | 150,000 |
| Voltage (kV) | 300 | 200 | 300 | 300 |
| Spherical Aberration (mm) | 2.7 | 2.7 | 2.7 | 0.01 |
| Electron Exposure ($e^-$/ Å$^2$) | 50 | 52 | 50 | 55 |
| Defocus range (mm) | -0.1 to -2.4 | -0.1 to -2.4 | -0.1 to -2.4 | -0.1 to -2.4 |
| Pixel size (Å, Physical/Digital) | 0.68 | 0.928 | 0.68 | 0.776 |
| Movies | 13,077 | 3826 | 13,077 | 18,000 |
| **Map Statistics and Post-Processing** | | | | |
| Symmetry imposed | C1 | C1 | C1 | C1 |
| Map Resolution (Å) | 3.12 | 3.33 | 3.03 | 3.01 |
| FSC threshold | 0.143 | 0.143 | 0.143 | 0.143 |
| **Model Composition** | | | | |
| Chains | 6 | 4 | 11 | 5 |
| Non-hydrogen atoms | 6314 | 10451 | 21695 | 12092 |
| Protein residues | 814 | 1342 | 1640 | 1131 |
| Ligands | CLF:2 | CLF:1<br>HCA:1<br>ICS:1 | CLF:1<br>HCA:1<br>ICS:1<br>ADP:3 | CLF:1<br>ICS:1 |
| Water | 0 | 0 | 0 | 0 |
| Bonds (RMSD)<br>Length (Å) (#>4s)<br>Angles (°) (#>4s) | <br>0.004 (0)<br>0.526 (0) | <br>0.004 (0)<br>0.738 (0) | <br>0.003 (0)<br>0.609 (0) | <br>0.010 (0)<br>0.582 (0) |
| **Validation** | | | | |
| Molprobity Score | 2.19 | 3.17 | 2.84 | 3.49 |
| Ramachandran plot | | | | |

| | | | | |
|---|---|---|---|---|
| Favored (%) | 94.66 | 87.54 | 90.46 | 88.89 |
| Allowed (%) | 5.34 | 12.39 | 9.36 | 11.11 |
| Outliers (%) | 0.00 | 0.08 | 0.19 | 0.00 |

**Supplementary Table 3.** Primers used in this study.

| Primer | Sequence (5′-3′) |
|---|---|
| P6.53 | GCCATTTTTCTTCTCACCGGGGAGCTGAGC |
| P6.64 | GCGCACCGTGGGCTTGTACTCGGTC |
| P6.65 | TCGCCTTCTTGACGAGTTCTTCTGAGCGGG |
| P7.3 | CAGCTAGCTAACGCGTATTAAAGG |
| P7.14 | CTGATGGTAGTCGAAGACG |
| P7.18 | CCTTCATAGTCATCCCTGTG |

**Supplementary Table 4.** Plasmids and Strains used in this study.

| Plasmid | Description | Reference |
|---|---|---|
| pAMG40 | *E. coli*-*Methanosarcina* shuttle vector for fosmid retrofitting, contains pC2A and λattB | Guss et al., *Archaea* 2008 |
| pDL238 | *M. acetivorans* CRISPR-Cas9 plasmid with no gRNA | Saini et al., *Scientific Reports* 2023 |
| pDL705 | pDL238 derivative with gRNA-*nifI2* (AGTGGTCCAGAAAATCATAG) and homology arms to add tandem Strep tag to N-terminus of *nifD* in *M. acetivorans* chromosome | This study |
| pDL707 | Cointegrate of pDL705 and pAMG40 | This study |
| pDL906 | pET28a derivative for expression of Strep-NifH from *M. acetivorans* in *E. coli* | This study |

| Strain | Description | Reference |
|---|---|---|
| *E. coli* DH5α | Commercial strain | |
| *E. coli* SufFeScient | BL21(DE3) but *zdh-3632*::*cat*, $^{-26}$ATA$^{-24}$ bp relative to *sufA* TSS changed to $^{-26}$TAT$^{-24}$ | Corless et. al., *J. Bac.* 2020 |
| *E. coli* WM4489 | *E. coli* DH10B derivative for pJK050 plasmid maintenance | Eliot et al., Chem. Biol. 2008 |
| *M. acetivorans* WWM73 | Δhpt::PmcrB-tetR-C31-int-attP | Guss et al., *Archaea* 2008 |
| *M. acetivorans* DJL70 | WWM73 *strep-nifD* | This study |

**Supplementary Table 5.** Molecular weights are particle distributions from Mass Photometry measurements.

| **Sample name** | **Observed Mass (kDa)** | **Error (kDa)** | **Area % under the curve** |
|---|---|---|---|
| PII-NifDK (30nM) | Peak 1 = *<br>Peak 2 = 847 | Peak 1 = *<br>Peak 2 = 158 | Peak 1 = 93.1<br>Peak 2 = 6.9 |
| PII-NifDK (10nM) | Peak 1 = 230 | Peak 1 = 105 | Peak 1 = 100 |
| PII-NifDK (10nM) +2OG+ATP | Peak 1 = 117<br>Peak 2 = 227 | Peak 1 = 21<br>Peak 2 = 104 | Peak 1 = 55<br>Peak 2 = 45 |
| PII-free NifDK (10nM) | Peak 1 = 101<br>Peak 2 = 169<br>Peak 3 = 239<br>Peak 4 = 437 | Peak 1 = 22<br>Peak 2 = 24<br>Peak 3 = 35<br>Peak 4 = 71 | Peak 1 = 22<br>Peak 2 = 33<br>Peak 3 = 43<br>Peak 4 = 2 |
